# A chromosome folding intermediate at the condensin-to-cohesin transition during telophase

**DOI:** 10.1101/678474

**Authors:** Kristin Abramo, Anne-Laure Valton, Sergey V. Venev, Hakan Ozadam, A. Nicole Fox, Job Dekker

**Author notes:** Correspondence: Job Dekker.

## Abstract

Chromosome folding is extensively modulated as cells progress through the cell cycle. During mitosis, condensin complexes fold chromosomes in helically arranged nested loop arrays. In interphase, the cohesin complex generates loops that can be stalled at CTCF sites leading to positioned loops and topologically associating domains (TADs), while a separate process of compartmentalization drives the spatial segregation of active and inactive chromatin domains. We used synchronized cell cultures to determine how the mitotic chromosome conformation is transformed into the interphase state. Using Hi-C, chromatin binding assays, and immunofluorescence we show that by telophase condensin-mediated loops are lost and a transient folding intermediate devoid of most loops forms. By late telophase, cohesin-mediated CTCF-CTCF loops and positions of TADs start to emerge rapidly. Compartment boundaries are also established in telophase, but long-range compartmentalization is a slow process and proceeds for several hours after cells enter G1. Our results reveal the kinetics and order of events by which the interphase chromosome state is formed and identify telophase as a critical transition between condensin and cohesin driven chromosome folding.

## Introduction

During interphase cohesin organizes chromosomes in loops, thought to be the result of a dynamic loop extrusion process ^1^. Loop extrusion can occur all along chromosomes but is blocked at CTCF sites leading to detectable loops between convergent CTCF sites ^2–7^ and the formation of topologically associating domains (TADs ^7–9^). At the same time long-range association of chromatin domains of similar state, within and between chromosomes, leads to a compartmentalized nuclear arrangement where heterochromatic and euchromatic segments of the genome are spatially segregated ^10^. Compartmentalization is likely driven by a process akin to microphase segregation and is mechanistically distinct from loop and TAD formation ^10–18^. The spatial arrangement of interphase chromosomes is closely correlated with gene expression patterns genome-wide, suggesting direct functional links between genome folding and gene regulation. The mechanistic connections, and causal relations, between chromosome organization and genome regulation are currently the topic of extensive studies ^19^.

During mitosis cohesin mostly dissociates from chromosome arms ^20, 21^ and condensin complexes re-fold chromosomes into helically arranged arrays of nested loops ^22–29^. This compaction facilitates accurate chromosome segregation during the metaphase-anaphase transition. Recently we described intermediate folding states through which cells interconvert the interphase organization into fully compacted mitotic chromosomes ^29^. During prophase cohesin-mediated loops, TADs and compartments disappear, and condensin II generates stochastically positioned arrays of loops. After nuclear envelope breakdown, during prometaphase, the combined action of condensins I and II generates nested loops and these start to follow a helical arrangement around a centrally located scaffold.

The kinetics and pathway of disassembly of the mitotic conformation and re-establishment of the interphase state as cells enter G1 are not known in detail. Previous studies have shown that condensin I loading, already high in metaphase, further increases during anaphase and then rapidly decreases, while condensin II colocalizes with chromatin throughout the cell cycle ^30^. Cohesin, mostly dissociated from chromatin during prophase and prometaphase ^20, 21^, re-associates with chromosomes during telophase and cytokinesis, as does CTCF ^20, 31, 32^.). However, it is not known how these events relate to modulation of chromosome conformation and whether distinct chromosome folding intermediates exist in the pathway towards a fully formed interphase nucleus. Here we used synchronized cell cultures to determine chromosome conformation, and chromatin association and dissociation of key chromatin architectural proteins such as condensins, cohesin and CTCF as cells exit mitosis and enter G1. We find that during telophase when condensin-dependent loops have dissolved, a transient chromosome conformation intermediate is formed that has properties that have previously been shown to be consistent with a fractal globule state and that is devoid of most loops. Along these chromosomes cohesin then re-loads, and cohesin mediated loops and TADs are established rapidly as cells progress through cytokinesis. Compartment domain boundaries are also detectable as early as telophase, but long-range, chromosome-wide association between distal compartment domains form with much slower kinetics and compartmentalization continues to strengthen until late G1.

## Results

### Synchronous entry into G1

HeLa S3 cells were arrested in prometaphase using our previously established protocol ^28^ (see methods). Briefly, cells were arrested in S-phase by addition of thymidine for 24 hours, released for 3 hours, and then arrested in prometaphase by the addition of nocodazole for 12 hours. FACS and microscopic inspection of cells showed that this procedure leads to cultures in which over 95% of cells are in prometaphase (Fig. 1a, Supplemental Fig 1, and see below). In order to determine how chromosome conformation changes as cells exit mitosis and enter G1, prometaphase arrested cells were released in fresh media (t = 0 hours) and aliquots were harvested at subsequent time points up to 12 hours after release from prometaphase. FACS analysis was used to determine the fraction of cells that had entered G1 based on their DNA content. We observed that about 50% of the cells had re-entered G1 between 3 and 4 hours after release from prometaphase arrest and that cells began to enter S phase after about 10 hours (Fig. 1a). The highest proportion of G1 cells was observed at 8 hours after release and data obtained at this time point is used as a G1 reference in this work. Replicate time courses yielded similar results with some variation in entry kinetics (Supplemental Fig. 2, 3).

**Fig. 1:**
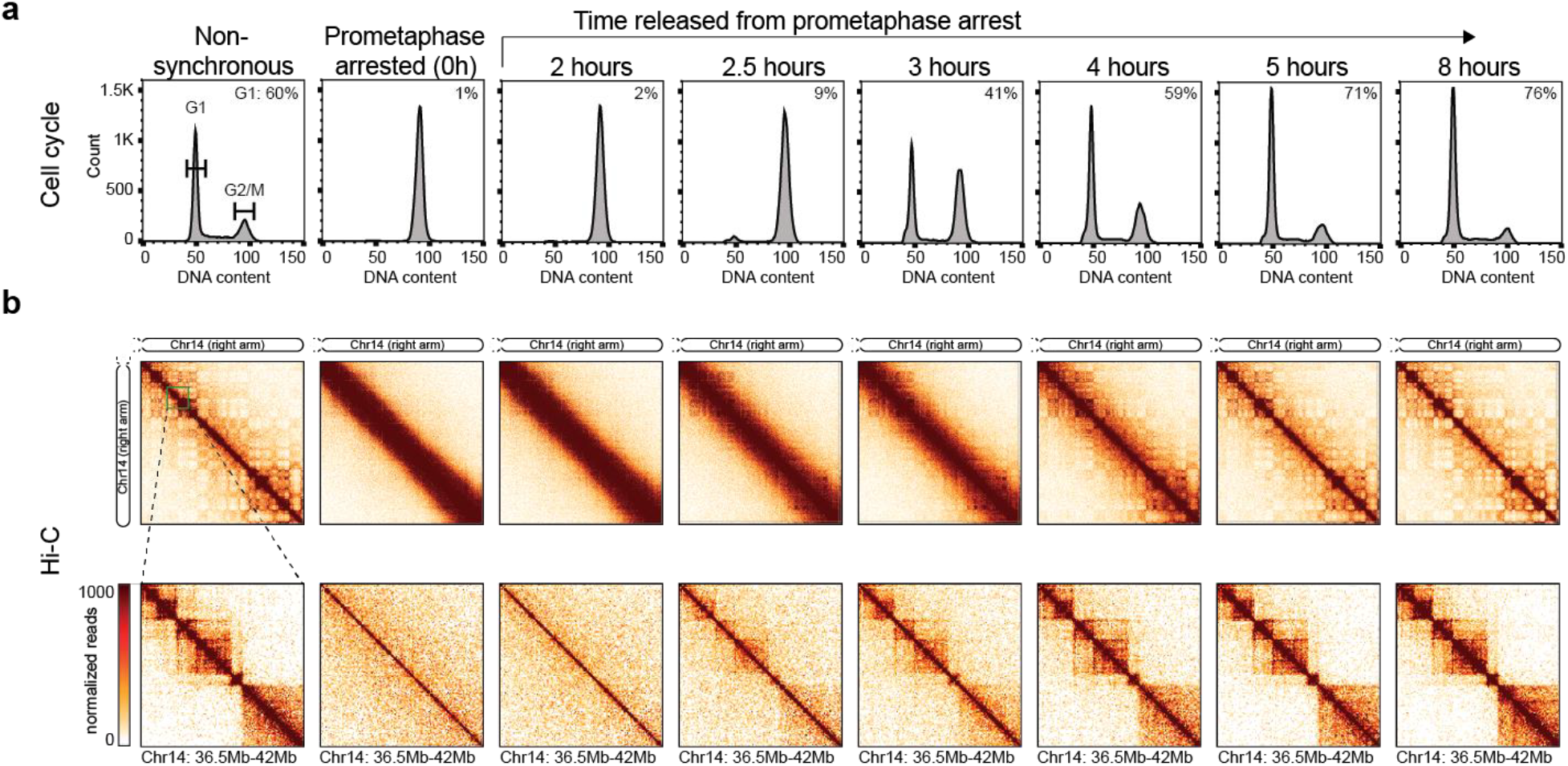
Hi-C analysis during mitotic exit and G1 entry. **a**, FACS analysis of nonsynchronous and prometaphase-arrested cultures and of cultures at different time points after release from prometaphase-arrest. Percentages in the upper right corner represent the percent of cells with a G1 DNA content. **b**, Hi-C interaction maps for nonsynchronous and prometaphase-arrested cultures and of cultures at different time points after release from prometaphase-arrest. Data for chromosome 14 are shown for two resolutions: 200 kb (top row) and 40 kb (bottom row).

### Chromosome conformational changes as cells enter G1

To assess chromosome conformation, we performed Hi-C on aliquots of cells taken at various time points after cells were released from prometaphase arrest (Fig. 1b). Hi-C chromatin interaction maps for cells in prometaphase reproduced previously identified features. First, the contact map is dominated by frequent interactions along the diagonal and the absence of locus specific features ^28^. When interaction frequencies (*P*) were plotted as a function of genomic distance (*s*) between loci, we observed the typical decay pattern observed for mitotic cells arrested with nocodazole (Supplementary Fig 4). *P*(*s*) initially decays slowly up to 10 Mb with an exponent close to −0.5, followed by a more rapid decay at larger distances. We note the presence of a weak shoulder in the decay plot around 10 Mb which might represent some remaining helicity that was shown to occur during earlier stages of prometaphase ^29^.

After release from prometaphase arrest, we observed a progressive gain in features of chromatin interaction maps normally seen in interphase and a reduced prominence of interactions across a broad diagonal typically seen in mitosis. First, visual inspection of the Hi-C interaction maps revealed the first emergence of short-range interphase chromatin features, such as TADs, as quickly as 2.5 hours after release from prometaphase arrest and these become more obvious over time (Fig. 1b, bottom row). Second, we observed the first appearance of a checker-board pattern of longer range interactions, reflecting the formation of A and B compartments, between 3 and 4 hours (Fig. 1b, top row). By 8 hours, the chromatin interaction maps and the shape of *P*(*s*) strongly resembled those obtained for nonsynchronous cell cultures (Supplementary Fig. 4) ^28^. We note that the checker-board at the 8 hour time point is sharper than that observed in a nonsynchronous culture, as expected for a pure G1 culture.

### Compartmentalization occurs slower than formation of TADs and loops

To confirm these visual observations, we quantified the presence and strength of specific features, such as TADs, loops, and compartments as they reform during mitotic exit and G1 re-entry. For these quantifications, we only used the set of structurally intact chromosomes in HeLa S3 cells as we did previously ^28^.

We used eigenvector decomposition to determine the positions of A and B compartments ^10^. In prometaphase-arrested cells, A and B compartments are absent, as expected (Supplemental Fig 5). Interestingly, by 3 hours release from prometaphase, PC1 detects the presence of A and B compartments, despite the fact that in Hi-C interaction maps, the checker-board pattern is weak (Fig. 1b, top row and Supplemental Fig 5a). For some chromosomes, PC3 corresponds to compartments at even earlier times (t = 2.75 hours) (Supplemental Fig. 5b). At subsequent time points, the amplitude of the PC1 track increases, which could be due to increased compartment strength as time progresses. To quantify compartment strength directly, we plotted interactions between loci arranged by their PC1 values (derived from the t = 8 hours Hi-C data) and obtained “saddle plots” ^14^ (Fig. 2a, top row). In these plots, interactions in the upper left corner represent interactions between B compartments and interactions in the lower right corner represent A-A interactions. The compartment strength is calculated as the ratio of homotypic (A-A and B-B) to heterotypic (A-B) interactions. The first appearance of preferred homotypic interactions is observed as early as 2.5 hours after release (Fig. 2a). These preferred interactions are initially weak, but gain strength during later time points. By ~5 hours after release, compartment strength is about 50% of the maximum strength we detect at 8 hours after prometaphase release.

**Fig. 2:**
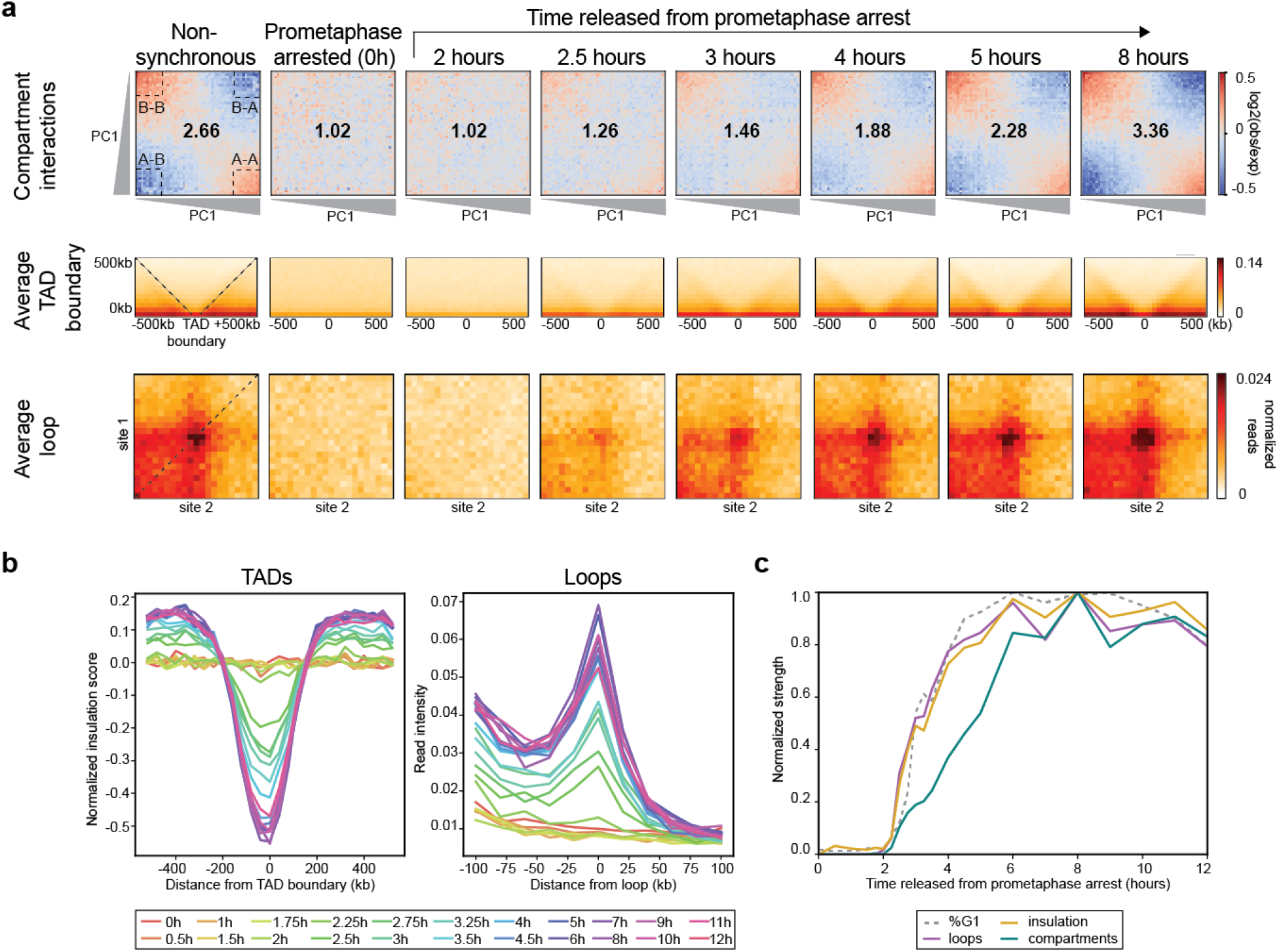
Kinetics of loop, TAD, and compartment formation. **a**, Top row: Saddle plots of Hi-C data binned at 200 kb resolution for nonsynchronous and prometaphase-arrested cultures and of cultures at different time points after release from prometaphase-arrest. Saddle plots were calculated using the PC1 obtained from the Hi-C data of the 8 hour time point. Numbers at the center of the heatmaps indicate compartment strength calculated as the ratio of AA+BB/AB using the mean values from dashed corner boxes. Middle row: Aggregate Hi-C data binned at 40 kb resolution at TAD boundaries identified from the Hi-C data of the 8 hour time point. The order of panels is the same as the top row. Dashed lines indicate the edges of the domains. Bottom row: Aggregate Hi-C data binned at 10 kb resolution at chromatin loops identified by Rao et al. ^2^. The order of panels is the same as the top row. **b**, Left: Average insulation profile across TAD boundaries for different time points. Right: Average Hi-C signals at and around looping interactions. Each line represents the signal from the lower left corner to the upper left corner of the loop aggregate heatmaps shown in panel a (dashed line). **c**, Normalized feature strength for TADs, loops, and compartments as a function of time after release from prometaphase. The strength for each of these features was set at 1 for the 8 hour time point. Dotted line indicates the fraction of cells in G1 at each time point, normalized to t = 8 hours.

We quantified A-A and B-B interaction frequencies separately as a function of time after release from prometaphase (Supplemental Fig. 6). We find that both compartment types form with similar kinetics (Supplemental Fig. 6c and 6f). Interestingly, when analyzed as a function of genomic distance between domains visual inspection of the saddle plots shows that B-B interactions are most prominent between loci separated up to 30 Mb, while A-A interactions are more prominent for loci separated by >30 Mb (Supplemental Fig. 6a and 6d). For compartment interactions up to 30 Mb, the kinetics of development of B-B interactions is faster than that of A-A interactions. For distances larger than 30 Mb, A-A interactions develop faster. These analyses reveal unanticipated complexities of compartmentalization.

Next, we quantified the appearance of domain boundaries, many of which define TADs. First, we calculated the insulation profiles along chromosomes ^33^. Insulation values represent the relative frequency of interaction across any locus. Minima in this profile represent domain boundaries ^34^. We aggregated Hi-C data at domain boundaries (Fig. 2a, middle row). In G1 cells, we observe a depletion of interactions across domain boundaries which is also illustrated by the average insulation profile across boundaries (Fig. 2b, left). In prometaphase, insulation at boundaries is absent, as observed before ^28^. As cells exit mitosis, we observe insulation at boundaries as soon as 2.5 hours after release from prometaphase. Insulation strength increases as time progresses and reaches 50% of maximum strength at ~3.5 hours after release. We are aware that some domain boundaries identified by insulation analysis represent compartment boundaries and not TAD boundaries. When analyzed separately, we find that compartment boundaries appear with similar kinetics as TAD boundaries (Supplemental Fig. 7). We conclude that both TAD and compartment domain boundaries are established around the same time, 2.5-3 hours after release from prometaphase.

Finally, we quantified the appearance of looping interactions. Rao et al. identified a set of 3,094 looping interactions in HeLa S3 cells, the large majority of which are interactions between CTCF sites that coincide with TAD boundaries, and 507 are on the structurally intact chromosomes in HeLa S3 cells ^2^. We aggregated Hi-C data at this set of looping interactions (Fig. 2a, bottom row). While such loops are readily detected in nonsynchronous cells, they are absent in prometaphase, as observed before ^35^. Loops reappear as soon as 2.5 hours after release and gain strength in the following hours. We calculated the strength of these loops by the Hi-C signal at the pairwise interaction divided by the Hi-C signals flanking it, illustrated in Fig. 2b. Loop strength reaches 50% of the maximum obtained over the time course after ~3.5 hours release from prometaphase.

To directly compare the kinetics with which TADs, loops, and compartments form, we plotted the strength of each feature at each time point as the percentage of its maximum during the time course (Fig. 2c). We observed that TADs and loops form with kinetics that are similar or slightly faster than the kinetics of G1 entry. In contrast, even though compartment identity is established relatively quickly (t = 2.5-3 hours), development and strengthening of long-range interactions between compartment domains, compartmentalization, continues for several hours with kinetics that are slower than that of cells entering G1. This temporal difference between TAD and loop formation, as compared to long-range inter-compartment interaction formation, might be related to the fact that these structures are formed by different mechanisms ^14–16^. Similar temporal kinetics of loop, TAD, and compartment formation was observed in independent time courses (Supplemental Fig 8).

### TADs and loops form prior to cytokinesis

TADs and loops appear somewhat earlier than cells starting to enter G1, but at later time points TAD and loop strength follows the accumulation of G1 cells closely. We reasoned that if the kinetics of TAD and loop formation is simply attributable to the kinetics of cells entering G1, then the observed Hi-C data at a given time point should be very similar to an appropriate mixture of a purely mitotic and purely G1 Hi-C dataset. To generate such mixtures, we randomly sampled reads from the prometaphase-arrested (t = 0 hour) and 8 hour released samples and mixed them according to the cell cycle distribution (percentage of cells in G1) of each sample to obtain a simulated time course of release from prometaphase (Fig. 3a). We then used the simulated time course datasets to perform the same analyses as described above to determine TAD, loop, and compartment strength (Fig. 3b-d).

**Fig. 3:**
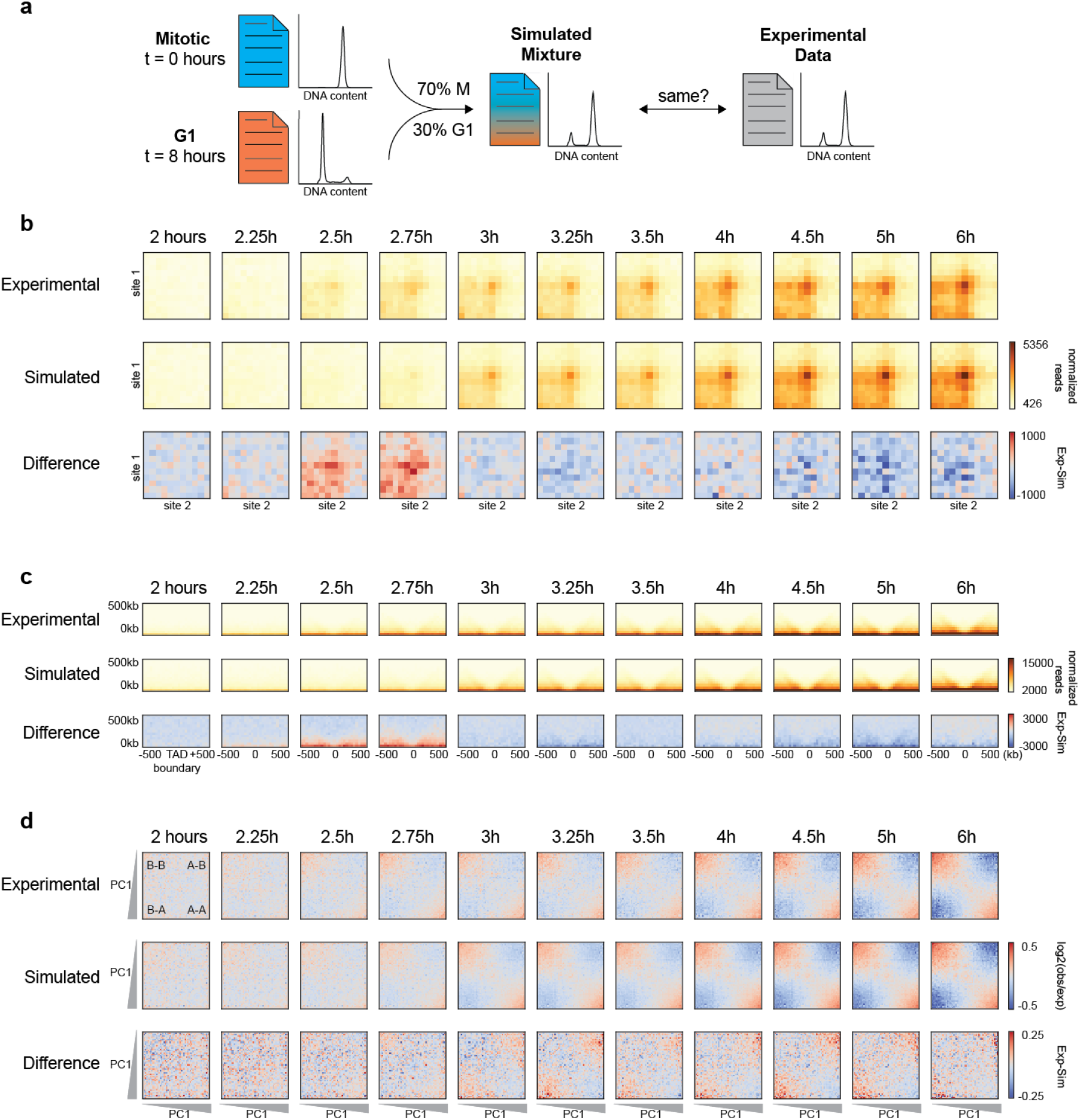
TADs and loops form quicker than expected, while compartmentalization occurs slower than expected. **a**, Schematic diagram of simulating Hi-C data based on the percentage of G1 cells at each time point. **b**, Aggregate Hi-C data binned at 20 kb resolution at chromatin loops at different time points. Top row: Experimental Hi-C data. Middle row: Simulated Hi-C data. Bottom row: The difference between experimental and simulated Hi-C data. Loops are more prominent in experimental Hi-C data than in the simulated data at t = 2.5 and t = 2.75 hours. This analysis included loops larger than 200 kb to avoid the strong signal at the diagonal of the interaction matrix. **c**, Aggregate Hi-C data binned at 40 kb resolution at TAD boundaries for different time points. Top row: Experimental Hi-C data. Middle row: Simulated Hi-C data. Bottom row: The difference between experimental and simulated Hi-C data. Insulation strength is stronger in experimental Hi-C data than in simulated Hi-C data at t = 2.5 and t = 2.75 hours. **d**, Saddle plots of Hi-C data binned at 200 kb resolution for different time points. Top row: Experimental Hi-C data. Middle row: Simulated Hi-C data. Bottom row: The difference between experimental and simulated Hi-C data. Saddle plots were calculated using the PC1 obtained from the experimental Hi-C data of the 8 hour time point. Compartmentalization is weaker in experimental Hi-C than in simulated Hi-C data as illustrated by the fact that A-B interactions are less depleted in the experimental data (upper right and lower left corner of saddle plots).

We first looked at the emergence of loops (Fig. 3b). In the experimental time course we observed loops at 2.5 hours after release from prometaphase. Interestingly, in the simulated time course loops appear later, at about 3 hours. To quantify the difference in loop strength between experimental and simulated data at each time point, we subtracted the average loop signal for these datasets at each time point. We find that at 2.5-2.75 hours after release, loop strength in the experimental data is greater than in the simulated data, indicating that the percentage of G1 cells is not predictive of loop strength at these early time points. We did not see a difference in the kinetics of loop formation for loops of different sizes, suggesting that loop extrusion is a relatively fast process (Supplemental Fig 9). Similarly, we quantified the appearance of insulation at TAD boundaries as a function of time in the experimental and simulated time course datasets (Fig. 3c). At 2.5 and 2.75 hours after release from prometaphase, TAD boundaries are more prominent in the experimental Hi-C data. Combined, this indicates that TADs and loops appear prior to cells entering G1. We used the same approach to determine how compartment strength relates to the fraction of cells in G1 (Fig. 3d). We quantified compartment strength using saddle plots, as above, and find that from 3 to 6 hours release, compartmentalization is weaker in the experimental Hi-C data as compared to the simulated Hi-C datasets: the simulated Hi-C data show less inter-compartment interactions (A-B) than the actual samples. This again illustrates that although compartment domain definition occurs as early as t = 2.5 hours, compartmentalization (the preferred long-range interaction between domains) is a relatively slow process that continues for several hours after cells have entered G1. Similar results were obtained with independent experimental and corresponding simulated time courses (Supplemental Fig. 10, 11).

### An intermediate folding state during mitotic exit

Properties of chromosome folding can be derived from *P*(*s*) plots. For example, *P*(*s*) plots for interphase and mitosis are distinct (Fig. 4a) and have been used to test models of chromosome folding ^1, 10, 28, 29, 36^. We calculated *P*(*s*) for Hi-C data obtained from cells at different times after release from prometaphase arrest. We observe a gradual transition over time from a mitotic *P*(*s*) shape to that of an interphase *P*(*s*) curve (Supplemental Fig. 4). The transitional shapes could be the result of a mixture of mitotic *P*(*s*) and interphase *P*(*s*) or could represent intermediate folding states. To distinguish these possibilities, we returned to our simulated mixtures of Hi-C data described above. We calculated *P*(*s*) for the simulated datasets and compared to experimental *P*(*s*) at each time point (Fig. 4a). For most of the time points, the simulated *P*(*s*) closely aligns with the experimental *P*(*s*) and the difference between the two plots is minor (Fig. 4a, bottom graphs). Interestingly, we observed relatively large differences when we compare simulated and experimental *P*(*s*) at 2.5 and 2.75 hours after release from prometaphase. This means that at those time points, the percentage of G1 cells (9% and 17%, respectively) does not explain the change in *P*(*s*), and indicates that at these times an intermediate chromatin folding state exists.

**Fig. 4:**
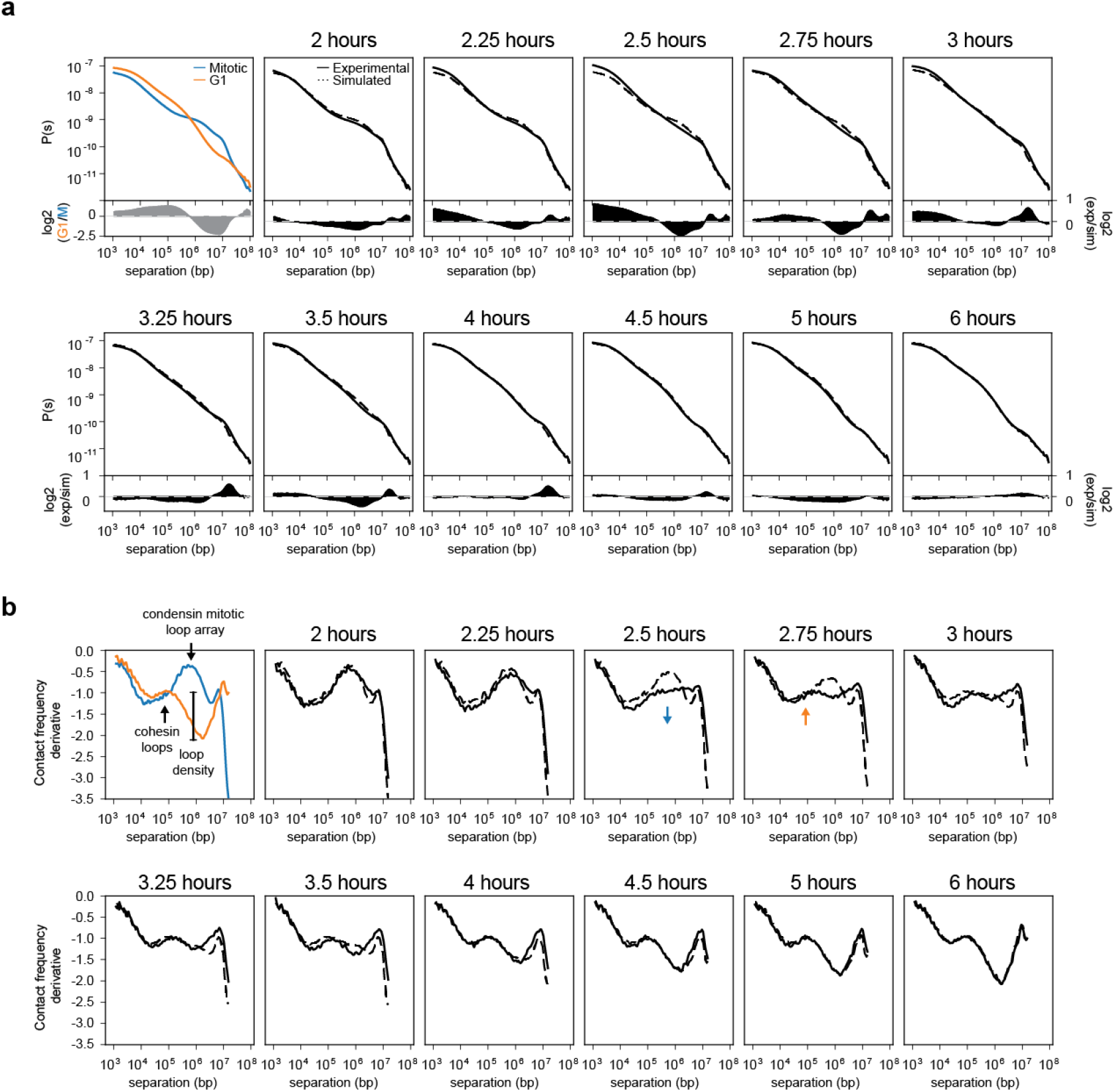
Formation of a transient folding intermediate. **a**, Contact frequency *(P*) versus genomic distance (*s*) for read normalized Hi-C datasets for experimental mitotic and G1 data (upper left, blue and orange lines, respectively) and experimental Hi-C data obtained from cells at different time points after release from prometaphase arrest (solid lines). Dotted lines are *P*(*s*) plots for simulated Hi-C datasets for corresponding time points. At the bottom of each *P*(*s*) plot, the difference between experimental and simulated *P*(*s*) is plotted for the different time points, except for the upper left plot which shows the difference *P*(*s*) for experimental G1 and mitotic cells. Note, that the difference plot for the upper left graph is on a different scale than all of the other difference plots. **b**, Derivative from *P*(*s*) plots shown in panel a. In the upper left graph, we indicate features that represent the condensin mitotic loop array and the cohesin loop size and density. The blue arrow indicates loss of the condensin-dependent mitotic loop array. The orange arrow indicates the initiation of the cohesin-dependent G1 loops.

To further explore this transition and the properties of this putative folding intermediate, we calculated the derivatives of *P*(*s*). Previous work has shown that the derivate of *P*(*s*) can reveal the average chromatin loop size and the density of loops along the chromosome ^29, 37, 38^. The derivative of *P*(*s*) for G1 cells shows a local maximum around 100 kb, indicating the average cohesin mediated loop size, followed by a relative deep minimum, indicating the linear density of chromatin loops (Fig. 4b). The derivative of *P*(*s*) for prometaphase cells shows a local maximum around several hundred kilobases. The interpretation of the derivative *P*(*s*) plots for densely packed loops arrays in mitosis is more complicated than the interpretation of interphase data. The maximum in the derivative of mitotic *P*(*s*) represents the condensin mediated loop array, but the size of the loops is likely smaller than the position of the maximum ^29^. Nonetheless, the derivate of *P*(*s*) for interphase and mitotic cells are highly distinct with the local maximum in interphase representing the cohesin-mediated loops and the local maximum in mitotic cells representing the presence of a densely packed condensin loop array. We compared derivatives of *P*(*s*) for simulated and experimental data across the time course (Fig. 4b, see Supplemental Fig. 10d and 11d for similar analyses of replicate time courses). We observe that experimental and simulated data are very similar for most time points. At 2.5 and 2.75 hours after release from prometaphase, however, the derivative of the experimental *P*(*s*) has a unique shape not observed at any time point in the simulated datasets, indicating it is not the result of a mixed population of mitotic and interphase cells. While for the simulated data evidence for a condensin loop array is still observable, the derivative of the experimental *P*(*s*) shows a relatively constant value of −1 for genomic distances ranging from 100 kb to 1 Mb. At subsequent time points, the local maximum around 100 kb becomes more prominent and the subsequent minimum becomes deeper indicating progressive cohesin loading and loop formation. We interpret this to mean that at t = 2.5 and t = 2.75 hours, there is a transient intermediate folding state in which the condensin loop array is largely disassembled and only some cohesin loops start to form. Similar results were obtained in three independent replicate time courses (Supplemental Fig. 10d, 11d). Inspection of the kinetics of the three time courses indicates that this intermediate conformation is transient, possibly as short as 15 minutes (Fig. 4b). Therefore, it is most reliably detected in time courses with many samples taken with short time intervals during the first hours after release from prometaphase arrest (e.g. compare time course 1 (Fig. 4b) to time course 2 (Supplemental Fig. 10d)).

### The transient intermediate folding state occurs at the anaphase-telophase transition

In order to better define the cell cycle state during which we observe the intermediate folding state characterized by a very low density of chromatin loops and no compartmentalization, we analyzed cells at different time points by microscopy. Specifically, we stained cells with DAPI to assess chromosome morphology and with antibodies against alpha-tubulin to detect spindle organization (Fig. 5a). Based on chromosome morphology and spindle organization, we classified cells (~74,000) into six different cell cycle phases: prometaphase, metaphase, early anaphase, late anaphase/early telophase, late telophase, and G1. We then determined the percentage of cells in each stage at different time points after release from prometaphase and generated cumulative plots (Fig. 5b). We observe that after 2.1 hours, 50% of the cells have entered metaphase and rapidly progress to anaphase (t = 2.2 hours). By 2.5 hours, 50% of the cells are at the anaphase to early telophase transition. Cells spend the next ~1.75 hours in telophase and cytokinesis and 50% of the cells have entered G1 after about 4 hours in this time course. From the timing of these events, we conclude that the transient intermediate folding state occurs during the anaphase-telophase transition between 2 and 3 hours after release from prometaphase.

**Fig. 5:**
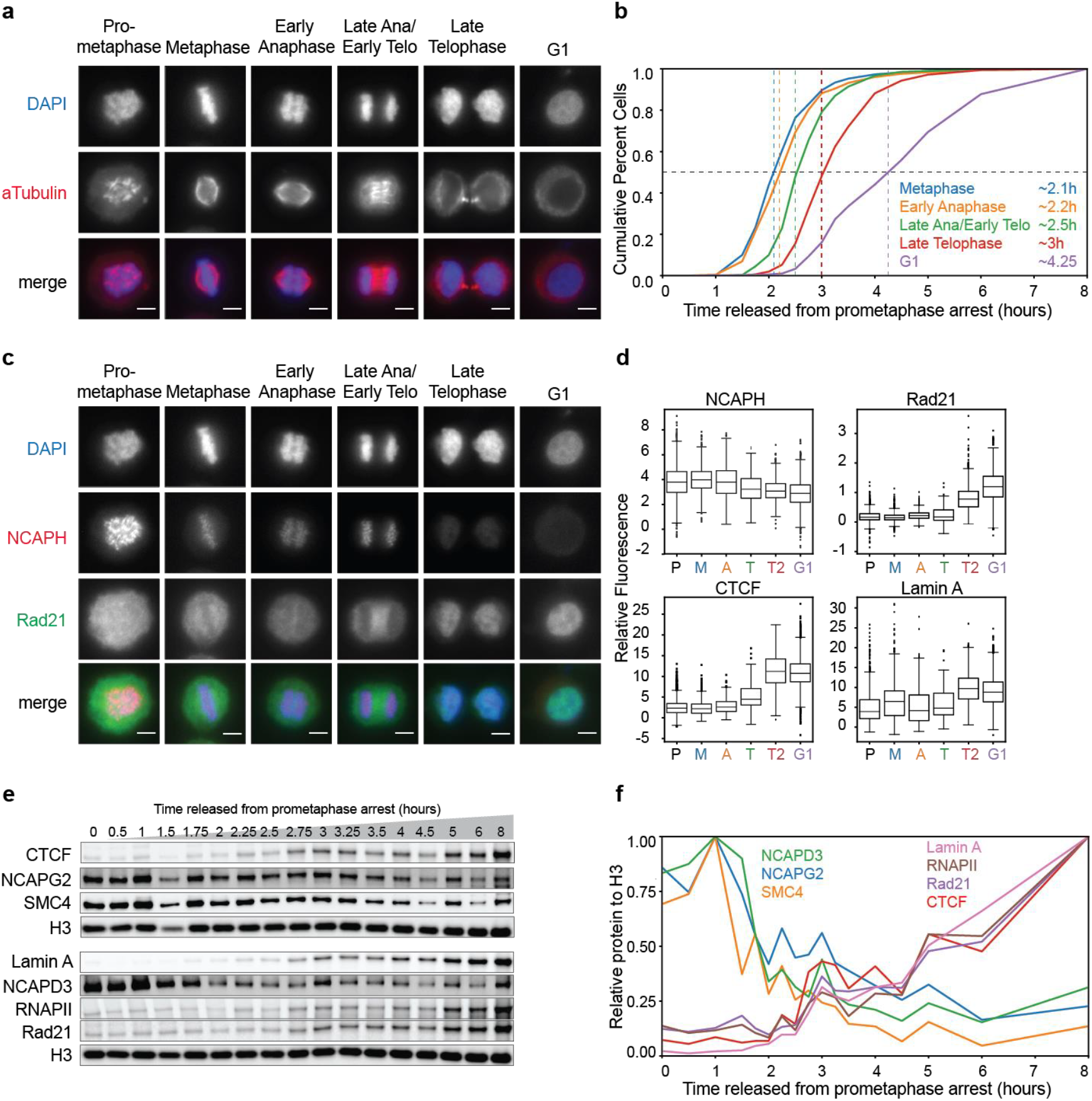
Chromatin association dynamics of condensins and cohesin during mitotic exit. **a**, Classification of cell cycle stages based on DAPI staining and alpha-tubulin organization. Prometaphase cells were defined as cells with condensed chromosomes and disrupted tubulin structure due to the microtubule inhibitor used for prometaphase arrest. Cells classified as metaphase had a single axis of DAPI staining with tubulin aligned on each side. Anaphase cells had tubulin on each side of the DAPI axis but must have had two distinct DAPI clusters representing the separation of two genomic copies. Late anaphase/early telophase classification was characterized by the presence of tubulin only between the two DAPI populations and no longer on the ends. When the tubulin signal was compressed between the two DAPI clusters, we classified those as late telophase and cytokinesis. Finally, all cells with decondensed chromatin and no nuclear tubulin were classified as G1 cells. Scale bar = 5 µm. **b**, Cumulative plots of cells at different cell cycle stages defined by imaging. **c**, Localization of NCAPH and Rad21 on chromosomes during different cell cycle stages. Scale bar = 5 µm. **d**, Quantification of NCAPH, Rad21 and CTCF colocalization with chromatin and Lamin A ring formation at different cell cycle stages (see Methods). P = prometaphase, M = metaphase, A = early anaphase, T = late anaphase/early telophase, T2 = late telophase, G1 = G1. **e**, Western blot analysis of chromatin-associated proteins purified from cells at different time points after release from prometaphase. **f**, Quantification of the western blot shown in panel e. Protein levels were normalized to Histone H3 levels from the same samples.

### Condensin unloading and cohesin loading occurs during the anaphase-telophase transition

The derivative of *P*(*s*) plots (Fig. 4b) combined with the cell cycle classification described above (Fig. 5b) indicate that the mitotic loop array is disassembled during late anaphase and early telophase. The mitotic loop array is generated by condensins I and II, while interphase loops and TADs are mediated by cohesin ^15, 16, 27, 29, 39^. Therefore, we set out to determine the kinetics with which condensins dissociate from chromosomes and cohesin associates with chromatin as cells exit mitosis. First, we analyzed condensin binding to chromosomes by microscopy. We have been unable to identify suitable antibodies for immunofluorescence detection of endogenous condensin I or II. Therefore, we generated a HeLa S3 cell line expressing the condensin I subunit NCAPH fused to dTomato. The kinetics of mitotic exit for this cell line are comparable, though slightly slower, to that of HeLa S3 cells (Supplemental Fig 12a). We again classified cells in different cell cycle stages based on chromosome morphology and spindle organization and analyzed condensin I binding (NCAPH-dTomato) and cohesin association (Rad21) (Fig. 5c-d). We find that condensin I is associated with chromosomes until late anaphase. By late telophase, most of the condensin I has dissociated. In contrast, very little cohesin is observed on chromosomes up until late anaphase, but is increasingly colocalized with chromatin during late telophase. In the same time course experiment, we also analyzed CTCF and Lamin A localization. We find that CTCF is not on chromosomes during prometaphase, consistent with our previous data ^35^. CTCF becomes colocalized with chromatin during late telophase with kinetics similar to that of cohesion (Fig 5d). Lamin A displays comparable chromatin colocalization dynamics.

Second, we determined chromatin association of these complexes directly by purifying chromatin-bound proteins followed by western blot analysis (Fig. 5e, left). Chromatin fractionation was used as an approach because many condensin and cohesin complexes associate with chromatin and generate loops in a sequence independent manner ^14, 29^, which is not readily detected using chromatin immunoprecipitation followed by DNA sequencing. We quantified the level of chromatin binding for proteins of interest from the western blot and normalized each to the Histone H3 level in the corresponding sample (Fig. 5e, right). We find that SMC4, a subunit of both condensin I and II, dissociates from chromatin rapidly during anaphase. Condensin II (NCAPG2, NCAPD3) showed very similar dissociation kinetics, as did condensin I (NCAPH-dTomato, Supplemental Fig. 12b). Cohesin (Rad21) started to associate with chromatin during anaphase-telophase and continued to load as cells entered and progressed through G1. Finally, we analyzed chromatin association of CTCF, Lamin A, and elongating RNAPII. They all show very similar binding kinetics as cohesin (Fig. 5e-f). The timing of chromatin association of cohesin and CTCF is consistent with earlier studies ^20, 31, 32^ and with more recent chromatin immunoprecipitation experiments that showed CTCF re-binding at CTCF sites during anaphase-telophase, followed rapidly by cohesin accumulation at those sites ^32, 40^.

We conclude that during late anaphase and into telophase, most condensin has dissociated from the chromosomes and both condensin and cohesin association with chromosomes is low. This is consistent with the interpretation of the Hi-C data based on the derivate of *P*(*s*) that at this time point there is a transient chromatin folding intermediate with no condensin-mediated loops and only a very low density of cohesin loops, and no long-range compartmentalization. As cells progress through late telophase and cytokinesis CTCF and cohesin increasingly load on chromosomes and this continues into G1.

## Discussion

We have determined how mitotic chromosome conformation is transformed into the interphase conformation (Fig 6). We identify late anaphase and telophase as a critical transition when the mitotic chromosome arrangement has been disassembled and features of the interphase conformation are becoming established. Hi-C, immunolocalization and chromatin binding assays show loss of condensin binding during late anaphase-early telophase while CTCF and cohesin start loading during telophase. At this critical juncture an intermediate chromosome conformation is detected that is characterized by the absence of most loops and no or very weak long-range inter-compartment interactions. This is a short-lived transient intermediate state. Subsequently during late telophase and cytokinesis CTCF and cohesin re-load, CTCF-CTCF loops and TAD boundaries are re-established as are compartment domains. While TADs and loops become more prominent rapidly with kinetics faster or equal to G1 entry, long-range compartmentalization occurs slower and continues to increase for several hours after cells have entered G1.

**Fig. 6:**
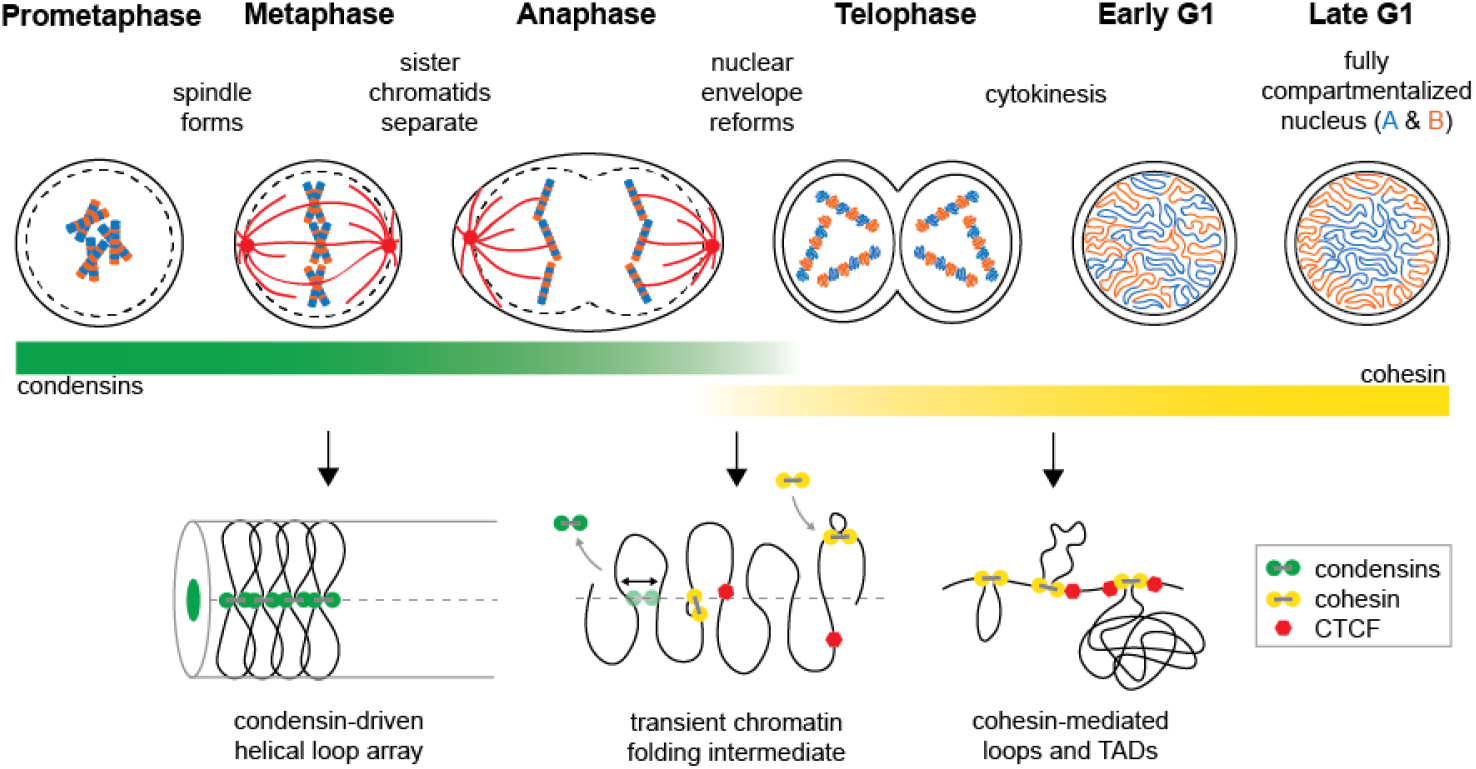
Cellular and chromosomal events as cells exit mitosis and enter G1. Top: Schematic diagrams indicate the cellular events including spindle formation, nuclear envelope reformation, and cytokinesis. Compartment type indicated by color: blue = A, orange = B. Bottom: Models of chromosome conformation during mitosis, telophase, and interphase. Green bar indicates abundance of condensins I and II on the chromatin at the corresponding cell cycle stages. Yellow bar indicates cohesin abundance on the chromatin at the corresponding cell cycle stages.

Our data show that key features that define the interphase state, loop anchors and domain boundaries (TADs and compartments) are defined prior to cells entering G1. The fact that TADs and loops form rapidly indicates that the process of loop extrusion is relatively fast, extruding loops of up to several hundreds of kb within 15-30 minutes, consistent with previous studies that showed that loop extrusion by SMC complexes is a fast process (1-2 Kb per second on naked DNA, ^41^, several Kb per minute during prophase ^29^. In contrast, long-range compartmentalization, i.e. the association of compartment domains that are typically separated by many megabases occurs more slowly during several hours in G1, even though their boundaries and identities are detectable much earlier (telophase). This supports the notion that compartmentalization is mechanistically distinct from TAD and loop formation. While the latter are formed by the active process of loop extrusion, the former has been proposed to be due to phase segregation that probably depends on passive diffusion ^11–13, 17, 18, 42^. A previous study also showed that compartmentalization occurs during early G1, although that work did not have time resolution during anaphase, telophase and cytokinesis ^43^. Our data are in line with very recent studies from the Blobel lab that independently found that domain and loop anchors are established prior to G1 entry while inter-compartment interactions develop slower ^40^. Further, our work suggests that B-B interactions are occurring at shorter genomic distances than A-A interactions (Supplemental Fig. 6). This may be related to the positioning of B domains at the lamina while A domains tend to be more central allowing for interactions between domains separated by larger distances. Additional studies are needed to further explore the relationship between compartmentalization and overall nuclear organization.

The formation of an intermediate folding state during telophase coincides with this condensin-to-cohesin transition. Hi-C data for this state shows that chromosomes are mostly devoid of loops and long-range compartmentalization is minimal. The exponent of *P*(*s*) for this intermediate fluctuates around −1 for loci separated by 100 kb up to several Mb. Interpretation of this feature is not straightforward. It could represent the fact that chromosomes are transitioning between two states, with the −1 exponent being the average of the two. Alternatively, and more interestingly, an exponent of ~-1 been proposed to correspond to a state that is similar to a fractal globule ^10, 44–46^. A key characteristic of fractal globule folding is that it represents a largely unentangled fiber. One speculative interpretation therefore is that during telophase chromosomes are topologically unentangled. How could this state be formed? One intriguing possibility is that this unentangled state is a remnant of the condensin-mediated mitotic loop array that is also not entangled. Continuous loop extrusion by condensin complexes, combined with topoisomerase II activity would lead to decatenation of adjacent loops ^47^. Indeed, loss of condensin at late mitotic stages leads to re-catenations of sister chromatids ^48^ suggesting continuous action of condensin may also be required to disentangle chromatin loops along each chromatid. Dissociation of condensin during anaphase would then leave a largely unentangled though still linearly arranged conformation. Subsequent cohesin loading would then initiate the formation of loops again. Although at this time the exact topological state of the telophase chromosomes is speculative, our results demonstrate that this transient state represents a key intermediate between the mitotic and interphase genome conformations. Future examination of the molecular and physical properties of this intermediate can not only reveal mechanisms by which cells build the interphase nucleus, but may also lead to better insights into the mitotic state from which it is derived.

## Acknowledgements

We thank Christina Baer from the UMass Imaging core for advice on imaging and help with the classification pipeline on CellProfiler. We thank members of the Dekker lab and Mirny lab for discussions. We acknowledge support from the National Institutes of Health Common Fund 4D Nucleome Program (DK107980), and the National Human Genome Research Institute (HG003143). J.D is an investigator of the Howard Hughes Medical Institute.

## Author contributions

K.A and J.D. conceived and designed the project. K.A and A.L.V. performed time courses. K.A. performed Hi-C, imaging, and chromatin-association experiments. A.N.F. generated the HeLa S3-NCAPH-dTomato cells and performed imaging experiments. K.A., S.V. and H.O. analyzed data. K.A. and J.D. wrote the manuscript.

## Supplementary Materials

Supplementary Figures 1-12

## Materials and Methods

### Cell Culture

HeLa S3 CCL-2.2 cells (ATCC CCL-2.2) and HeLaS3-NCAPH-dTomato cells (see below) were cultured in DMEM, high glucose, GlutaMAX™ Supplement with pyruvate (Gibco 10569010) with 10% fetal bovine serum (Gibco 16000044) and 1% PenStrep (Gibco 15140) at 37°C in 5% CO_2_.

### Creation of Stable HeLaS3-NCAPH-dTomato Cell Line

We used pSpCas9(BB)-2A-Puro (PX459) V2.0 [a gift from Feng Zhang (Addgene plasmid # 62988; http://n2t.net/addgene:62988; RRID:Addgene_62988)] to construct CRISPR/Cas vectors according to the protocol of Ran et al. ^49^. gRNAs are listed in Table 1.

**Table 1.**
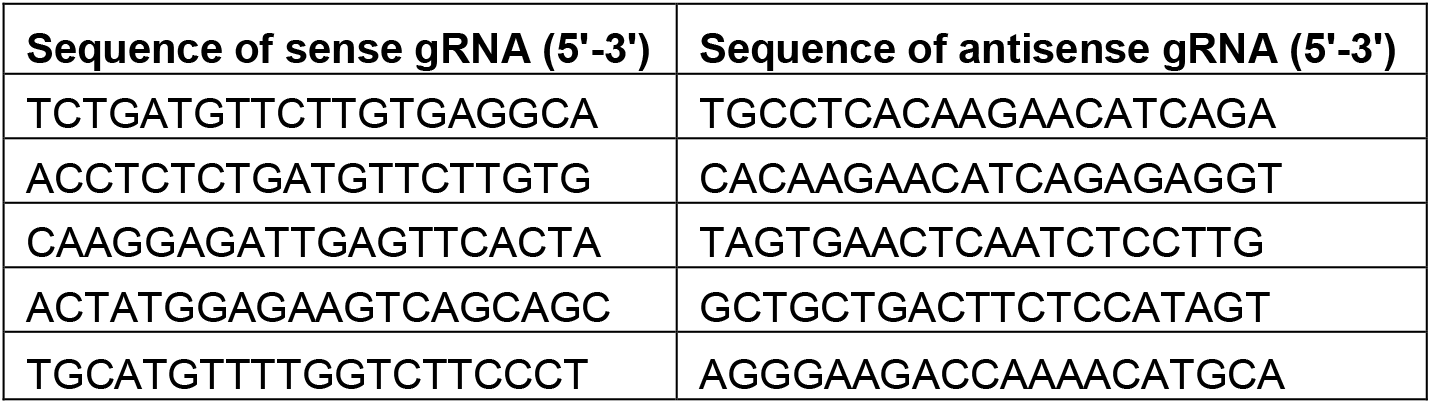
gRNA sequences for genome editing

To construct donor plasmids for C-terminal integration of dTomato, plasmids were based on pUC19 and constructed using synthesized DNA and homology arms generated by PCR (primers listed in Table 2). Template DNA (genomic DNA from HeLa S3 cells) was amplified using Q5 High-Fidelity DNA Polymerase (New England Biolabs) to generate NCAPH homology arms. gBlock containing dTomato and Blasticidin resistance was synthesized by Integrated DNA Technologies (IDT) (sequence in Table 3). Homology arms and gBlocks were cloned into pUC19 by Gibson assembly, using NEBuilder® HiFi DNA Assembly Master Mix (NEB).

**Table 2.**
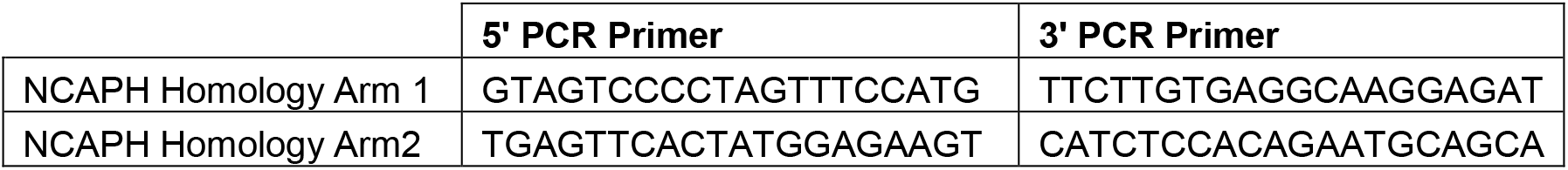
PCR primers for NCAPH homology arms

**Table 3.**
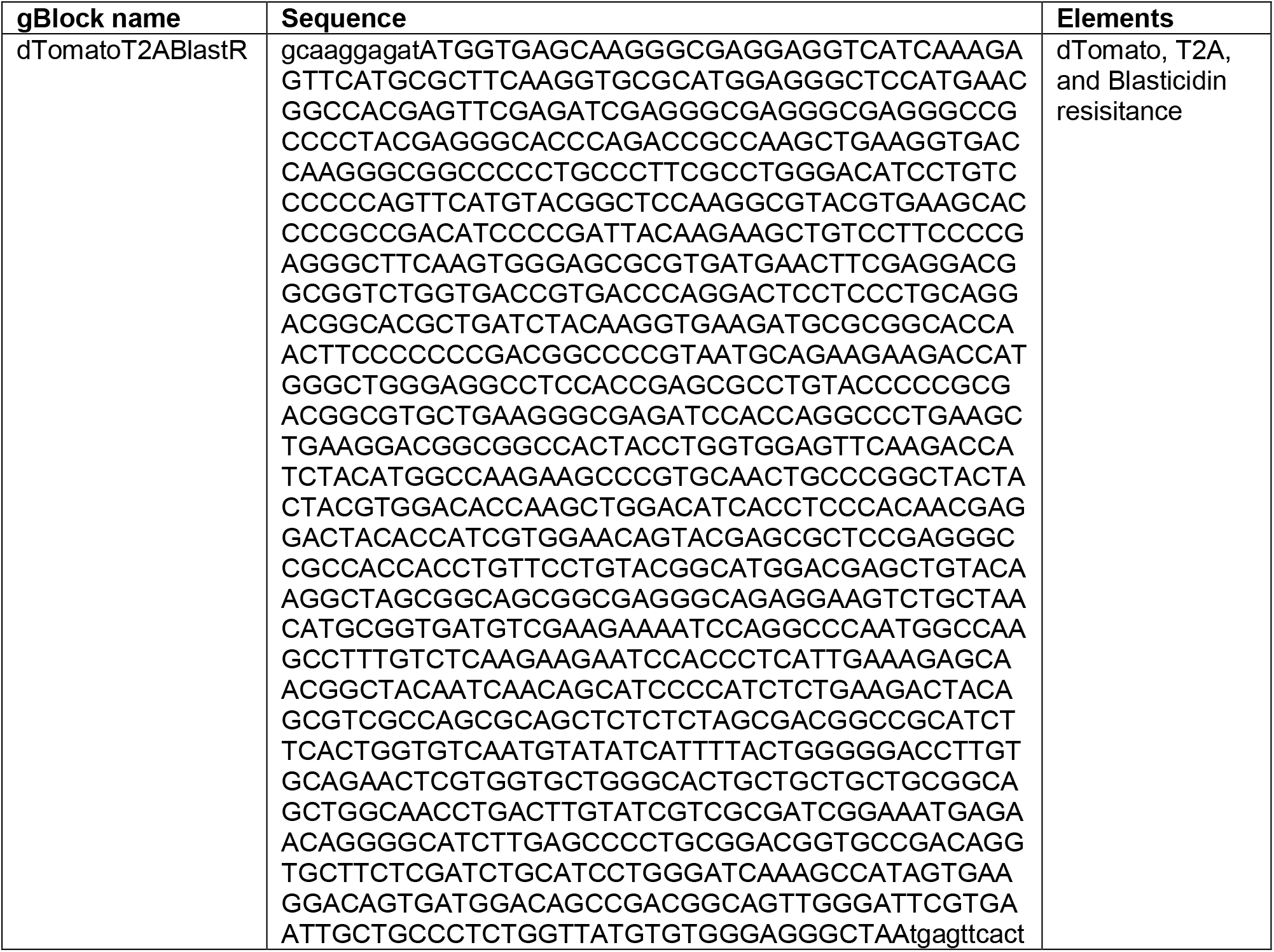
gBlock Gene Fragment

To generate stable cell lines, 5 × 10^6^ cells were electroporated with gRNAs and donor plasmid. 24 hours after electroporation, 1 μg/ml puromycin was added. Two days later, 1 ug/mL blasticidin was added for NCAPH-dTomato selection. After 5 days, colonies were picked for further selection in a 96-well plate.

### Mitotic Synchronization

All prometaphase synchronization of cells were done by (1) single thymidine treatment to arrest cells in S phase, (2) release into standard media to allow cell recovery and entry into late S, and (3) nocodazole treatment to arrest cells in prometaphase. On Day 1, cells were plated at 4 × 10^6^ cells / 15 cm plate in media containing 2mM thymidine (Sigma T1895). After 24 hours, cells were washed with 1X PBS (Gibco 14190144) and standard media was added back to plates for 3 hours. Cells were then treated with media containing 100 ng/mL nocodazole (Sigma M1404) for 12 hours. Floating mitotic cells were collected and washed in 1X PBS.

### Mitotic Release Timecourse

For prometaphase samples, washed mitotic cells were immediately prepared for downstream analysis. Remaining samples were re-cultured in standard media for synchronous release into G1 and collected at indicated times. For early time points, both floating and adherent re-cultured cells were collected for analysis. After 5 hours release from nocodazole, only adherent cells were collected.

Approximately 5 × 10^6^ cells at each time point were fixed in 1% Formaldehyde (Fisher BP531-25) diluted in serum-free DMEM for Hi-C analysis as described in Belaghzal et al. ^50^. For cell cycle analysis, approximately 1 × 10^6^ cells at each time point were fixed in 86% cold ethanol (Fisher 04-355-222) and stored at −20°C. For chromatin association protein analysis, approximately 5 × 10^6^ cells at each time point were pelleted, flash frozen, and stored at −80°C. Additional samples were collected for fluorescent microscopy. Floating mitotic cells were resuspended in 1.5 mL 4% PFA (EMS 15710) (diluted in 1X PBS), transferred onto a Poly-L-lysine-coated coverslip (Sigma P8920) in a 6 well plate, and spun at 1500xg for 15 min. Cells adherent to coverslips at later time points were fixed in 4% PFA for 15 minutes at 20°C. All coverslips were washed 3X in 1X PBS and stored in 1X PBS at 4°C.

### Cell Cycle Analysis

Fixed cells were washed in 1X PBS then resuspended in PBS containing 0.1% NP-40 (MP Biomedicals 0219859680), 0.5 mg/mL RNase A (Roche 10109169001) and 50 ug/mL propidium iodide (Thermo P1304MP). Samples were incubated at 20°C for 30 minutes then analyzed via LSR II or MACSQuant VYB flow cytometry. Data was analyzed using FlowJo v3. Viability gates using forward and side scatter were set on the nonsynchronous sample and applied to all samples within the set. DNA content was plotted as a histogram of the red channel. G1, S, and G2/M gates were set on nonsynchronous sample and applied to all samples within the set to get percentage of cells in each state throughout the time course release from prometaphase arrest. Values plotted for kinetics of G1 entry were normalized such that the maximum number of G1 cells = 1.

### Hi-C Protocol

Hi-C was performed as described in Belaghzal et al. ^50^. Briefly, cross-linked cell culture samples were lysed then digested with DpnII at 37°C overnight. Next, the DNA overhanging ends were filled with biotin-14-dATP at 23°C for 4 hours and ligated with T4 DNA ligase at 16°C for 4 hours. DNA was then treated with proteinase K at 65°C overnight to remove crosslinked proteins. Ligation products were purified, fragmented by sonication to an average size of 200 bp, and size selected to fragments 100 – 350 bp. We then performed end repair and dA-tailing and selectively purified biotin tagged DNA using streptavidin beads. Illumina TruSeq adaptors were added to form the final Hi-C ligation products, samples were amplified and PCR primers were removed. Hi-C libraries were then sequenced by PE50 bases on an Illumina HiSeq4000.

### Hi-C Data Processing

Hi-C PE50 fastq sequencing files were mapped to hg19 human reference genome using *distiller-nf* mapping pipeline (https://github.com/mirnylab/distiller-nf). In brief, bwa mem was used to map fastq pairs in a single-side regime (-SP). Aligned reads were classified and deduplicated using *pairtools* (https://github.com/mirnylab/pairtools), such that uniquely mapped and rescued pairs were retained and duplicate pairs (identical positions and strand orientations) were removed. We refer to such filtered reads as valid pairs. Valid pairs were binned into contact matrices at 10 kb, 20 kb, 40 kb, and 200 kb resolutions using *cooler* ^51^. Iterative balancing procedure ^52^ was applied to all matrices, ignoring the first 2 diagonals to avoid short-range ligation artifacts at a given resolution, and bins with low coverage were removed using MADmax filter with default parameters. Resultant “.cool” contact matrices were used in downstream analyses using *cooltools* (https://github.com/mirnylab/cooltools). For downstream analyses using *cworld* (https://github.com/dekkerlab/cworld-dekker), contact matrices were converted to “.matrix” using *cooltools dump_cworld*. For visualization of contact matrices (as in Fig. 1),. matrix files were scaled to 100 × 10^6^ reads using *cworld scaleMatrix*.

### Contact probability (*P*(*s*)) plots & derivatives

Cis reads from the valid pairs files were used to calculate the contact frequency (*P*) as a function of genomic separation (*s*) (adapted from *cooltools*). All *P*(*s*) curves were normalized for the total number of valid interactions in each data set. Corresponding derivative plots were made from each *P*(*s*) plot.

### Compartment analysis

Compartment boundaries were identified in cis using eigen vector decomposition on 200 kb binned data with *cooltools call-compartments* function. A and B compartment identities were assigned by gene density tracks such that the more gene-dense regions were labeled A compartments, and the PC1 sign was positive. Change in compartment type, therefore, occurs at locations where the value of PC1 changes sign. Compartment boundaries were defined at these locations, except for when the sign change occurred within 400 kb of another sign change.

To measure compartmentalization strength, we calculated observed/expected Hi-C matrices for 200 kb binned data, correcting for average distance decay as observed in the *P*(*s*) plots (*cooltools compute-expected*). We then arranged observed/expected matrix bins according to their PC1 values of the replicate 1 Hi-C dataset from cells released from prometaphase for 8 hours. We aggregated the ordered matrices for each chromosome within a dataset then divided the aggregate matrix into 50 bins and plotted, yielding a “saddle plot” (*cooltools compute-saddle*). Strength of compartmentalization was defined as the ratio of (A-A + B-B) / (A-B + B-A) interactions. Values used for this ratio were determined by calculating the mean value of the 10 bins in each corner of the saddle plot. Values plotted for kinetics of compartment formation were normalized such that strength = 0 in prometaphase cells and the maximum value = 1.

In order to observe compartmentalization at different genomic ranges, we extracted observed/expected Hi-C data at specific distances (0-12 Mb, 12-24 Mb, 24-36 Mb, 36-48 Mb, and 48-60 Mb) and made saddle plots. Since less data was used as input for each saddle plot, data was split into 20 bins instead of 50. Overall compartmentalization strength was calculated similar to above except using the mean value of the 9 bins in each corner of the saddle plot. Compartmentalization of individual compartment types was defined as the ratio of A-A / A-B or B-B / A-B, where these values were determined by calculating the mean value of the 9 bins in the specified corner of the saddle plot. All values were normalized and plotted for kinetics the same as above.

### TAD analysis

Domain boundaries were identified using insulation analysis on 40 kb binned data with *cworld matrix2insulation* and locating the minima in each profile (--is 520 kb --ids 320 kb). Domain boundaries were classified as compartment boundaries if they overlapped with the compartment boundaries defined above. All other domain boundaries were assumed to be TAD boundaries. To measure TAD boundary formation, we aggregated 40 kb binned Hi-C data at domain boundaries identified from the replicate 1 Hi-C dataset from cells released from prometaphase for 8 hours (*cworld elementPileUp*). Strength of TAD boundary formation was defined as the depletion of interactions across the boundary pileup, i.e. insulation. Boundary strength was calculated by measuring the average interaction of domain boundaries with regions 40-500 kb away (center vertical bin of boundary pileup) and subtracting that value from the average signal in regions immediately flanking the domain boundary (all bins left and right of domain boundary). All calculations were made after removing the bin closest to the diagonal. Values plotted for kinetics of TAD formation were normalized such that strength = 0 in prometaphase cells and the maximum value = 1.

### Loop analysis

We used a previously identified set of HeLa S3 looping interactions for this analysis ^2^. This set contains 3,094 total loops and 507 looping interactions are on the structurally intact chromosomes of HeLa S3 cells ^28^. To visualize looping interactions observed, we aggregated 10 or 20 kb binned data at loops larger than 200 kb to avoid the strong signal at the diagonal of the interaction matrix (*cworld interactionPileUp)*.

Strength of loop formation was defined as the enrichment of signal at the looping interactions (center 3×3 pixels at loop position 20 kb binned data) compared to the flanking regions. Strength was calculated by averaging the signal at the looping interaction and subtracting the average signal outside. Values plotted for kinetics of loop formation were normalized such that strength = 0 in prometaphase cells and the maximum value = 1.

In order to observe formation of looping interactions at all loops sizes, we aggregated observed/expected Hi-C matrices for 20 kb binned Hi-C data at sites of looping interactions. Using the observed/expected matrices corrects for distance decay and removes the overwhelming signal close to the diagonal, allowing us to observe smaller loops than in the observed Hi-C matrices.

### Simulated Hi-C mixture datasets

We generated simulated Hi-C datasets for each replicate time course experiment. For each replicate the following protocol was used to randomly mix reads from prometaphase Hi-C datasets (t = 0 hours) with random Hi-C data reads from the sample having the highest percentage of G1 cells in the respective time course (t = 8 hours for replicate 1 and 2, t = 6 hours for replicate 3). Mixing ratios were determined based on cell cycle analysis of the same time course replicate, such that x% prometaphase reads + 1-x% G1 reads was representative of the experimental FACs profile observed at each time point.

First, in order to properly compare samples, all valid pair files within a single Hi-C timecourse dataset were randomly down-sampled to the lowest number of uniquely mapped reads within that timecourse dataset. Next, the down-sampled valid pairs for experimental prometaphase (t = 0 hours) and experimental G1 (t = 6 or 8 hours) were randomly sampled to yield the correct ratio of experimental cells at each time point and the same number of total reads as the down-sampled valid pairs files. This step was repeated 25 times, resulting in 25 simulated valid pairs files with the same number of reads for each time point in each replicate. *P*(*s*) plots for simulated Hi-C data represent the average *P*(*s*) for 25 replicate valid pair simulations. For all other analyses, valid pairs files were binned and balanced (as above) into “.cool” contact matrices and the 25 replicates from the same simulated ratios were combined using *cooler merge*.

### Microscopy—staining & analysis

#### Immunofluorescence staining

Immunofluorescence staining was performed at room temperature. Fixed cells were permeabilized with 0.1% triton (Sigma T8787) in 1X PBS for 10 minutes. Cells were blocked with 3% BSA (Sigma A7906) in 0.1% triton/PBS for 1 hour. Cells were incubated with primary antibody diluted in the blocking buffer for 2 hours [alpha-tubulin mouse mAb (1:5000, Sigma T6199), Rad21 rabbit pAb (1:1000, ab154769), Lamin A rabbit pAb (1:1000, ab26300), CTCF rabbit pAb (1:800, Cell Signaling 2899)]. Cells were washed with 0.1% triton/PBS 3 × 5 minutes. Cells were incubated with secondary antibody diluted in the blocking buffer for 1 hour in the dark [Alexa Fluor 647 AffiniPure goat anti-mouse IgG (H+L) (1:100, Jackson 115-605-062), goat anti-rabbit IgG H&L Alexa Fluor 488 (1:1000, ab15007), goat anti-mouse IgG H&L Alex Fluor 568 (1:1000, ab175473)]. Cells were washed with 0.1% triton/PBS 1 × 5 minutes and then washed with 1X PBS 3 × 5 minutes. Coverslips were mounted to slides using ProLong Diamond Antifade Mountant with DAPI (Invitrogen P36962). For image acquisition, we used a Nikon Eclipse Ti microscope. Imaging was performed using an Apo TIRF, N.A. 1.49, 60x oil immersion objective (Nikon) and a Zyla sCMOS camera (Andor). Images were acquired using NIS-Elements 4.4.

#### Cell Cycle Classification

A separate training set of over 1000 individual HeLa S3 cells stained with DAPI and alpha-tubulin was used to set the classification parameters in Cell Profiler 3.1.8 and Cell Profiler Analyst 2.2.1 ^53–55^. Nd2 files from above were split into individual tiffs by channel. DAPI and alpha-tubulin intensity, shape, and texture were measured for each cell, and cells were classified into either prometaphase, metaphase, early anaphase, late anaphase/early telophase, late telophase, G1, or negative populations. Cell cycle classifications were confirmed by visual inspection. A total of ~74,000 cells were classified in this study.

#### Protein Localization

Cell Profiler was also used to measure the localization of NCAPH-dTomato, Rad21, CTCF, and Lamin A. For each cell, we identified primary objects in the DAPI channel (‘DNA’), we used propagation to look for secondary objects in the alpha-tubulin channel (‘tubulin’), and finally we created a tertiary object as the region between the primary and secondary objects (‘cytoplasm’). We calculated enrichment of NCAPH, Rad21, and CTCF co-localizing with the chromatin by measuring the mean fluorescence intensity (MFI) of each protein overlapping with the ‘DNA’ object and subtracting the MFI of each protein overlapping with the ‘cytoplasm’ object.

To measure the formation of a lamin ring, we shrunk the ‘DNA’ object and subtracted this region from the ‘DNA’ original object to create a new object (‘lamin’) at the inside edge of the ‘DNA’ where we observed lamin ring presence in nonsynchronous cells. Next, we expanded the ‘DNA’ object and subtracted the original ‘DNA’ object to create a new object (‘LamCyto’) just outside of the ‘lamin’ object. We were able to quantify the presence of a lamin ring by subtracting the MFI of lamin fluorescence in ‘LamCyto’ region from the MFI of lamin in the ‘lamin’ region. This enriched for the signal of a lamin ring, therefore, higher values correlated with the presence of a lamin ring structure at the edge of the chromatin.

### Chromatin association

#### Fractionation protocol

Flash-frozen cell pellets from each time point of the mitotic release time course were thawed and resuspended with lysis buffer (50 mM Tris-HCl, pH 8.0, 100 mM NaCl, 1% NP-40, 1 mM DTT, 1X Halt protease inhibitor cocktail (Thermo 78430)). Samples in lysis buffer were incubated on ice for 20 minutes and then spun at 13,000 × g for 10 minutes at 4°C. The supernatant (cytoplasmic fraction) was collected and the pellet was resuspended in nuclei buffer (10 mM PIPES, pH 7.4, 10 mM KCl, 2 mM MgCl_2_, 0.1% NP-40, 1 mM DTT, 1X protease inhibitor) with 0.25% triton. Samples were incubated on ice for 10 minutes and then spun at 10,000 × g for 5 minutes at 4°C. The supernatant (nucleoplasmic fraction) was collected and the pellet (chromatin fraction) was resuspended in nuclei buffer with 20% glycerol. The chromatin fraction was then sonicated to shear the DNA using a Covaris instrument with the following parameters: 10% duty cycle, intensity 5, 200 cycles/burst, frequency sweeping, continuous degassing, 240 second process time, 4 cycles. Final chromatin-bound protein samples were stored at −20°C.

#### Western Blots

The volume for approximately the same number of cells for each sample across the mitotic release time course was loaded in each lane of a 4-12% bis-tris protein gel (Biorad 3450125) and separated in 1X MES running buffer (Biorad 1610789). Proteins were transferred to nitrocellulose membranes (Bio-Rad 1620112) at 30 V for 1.5 hours in 1X transfer buffer (Thermo 35040). Membranes were blocked with 4% milk in PBS-T (1X PBS + 0.1% tween) for 1 hour at room temperature. Membranes were then incubated with specified primary antibody diluted 1:1000 in 4% milk/PBS-T overnight at 4°C [Histone H3 (ab1791), Rad21 (ab154769), RFP (cross-reacts with dTomato for NCAPH-dTomato, Rockland 600-401-379), SMC2 (ab10412), SMC4 (ab17958), NCAPD3 (ab70349), NCAPG2 (ab70350), Lamin A (ab26300), CTCF (Cell Signaling 2899), RNA polymerase II CTD repeat phospho S2 (ab5095)]. Membranes were washed with PBS-T 3 × 10 minutes at room temperature, then incubated with secondary antibody (anti-rabbit IgG HRP-linked, Cell Signaling 7074) diluted 1:4000 in 4% milk/PBS-T for 2 hours at room temperature. Membranes were washed with PBS-T 3 × 10 minutes. Membranes were developed and imaged using SuperSignal West Dura Extended Duration Substrate (Thermo 34076) and Bio-Rad ChemiDoc.

#### Quantification

Band intensity for each protein was quantified using Image Lab 5.2.1. Intensities for each lane were normalized by background intensity of an equal sized area in the same lane. All protein quantifications were normalized to the Histone H3 levels for the same time course samples.

