## Supplemental Figures and Tables for "A chromosome folding intermediate at the condensin-to-cohesin transition during telophase"

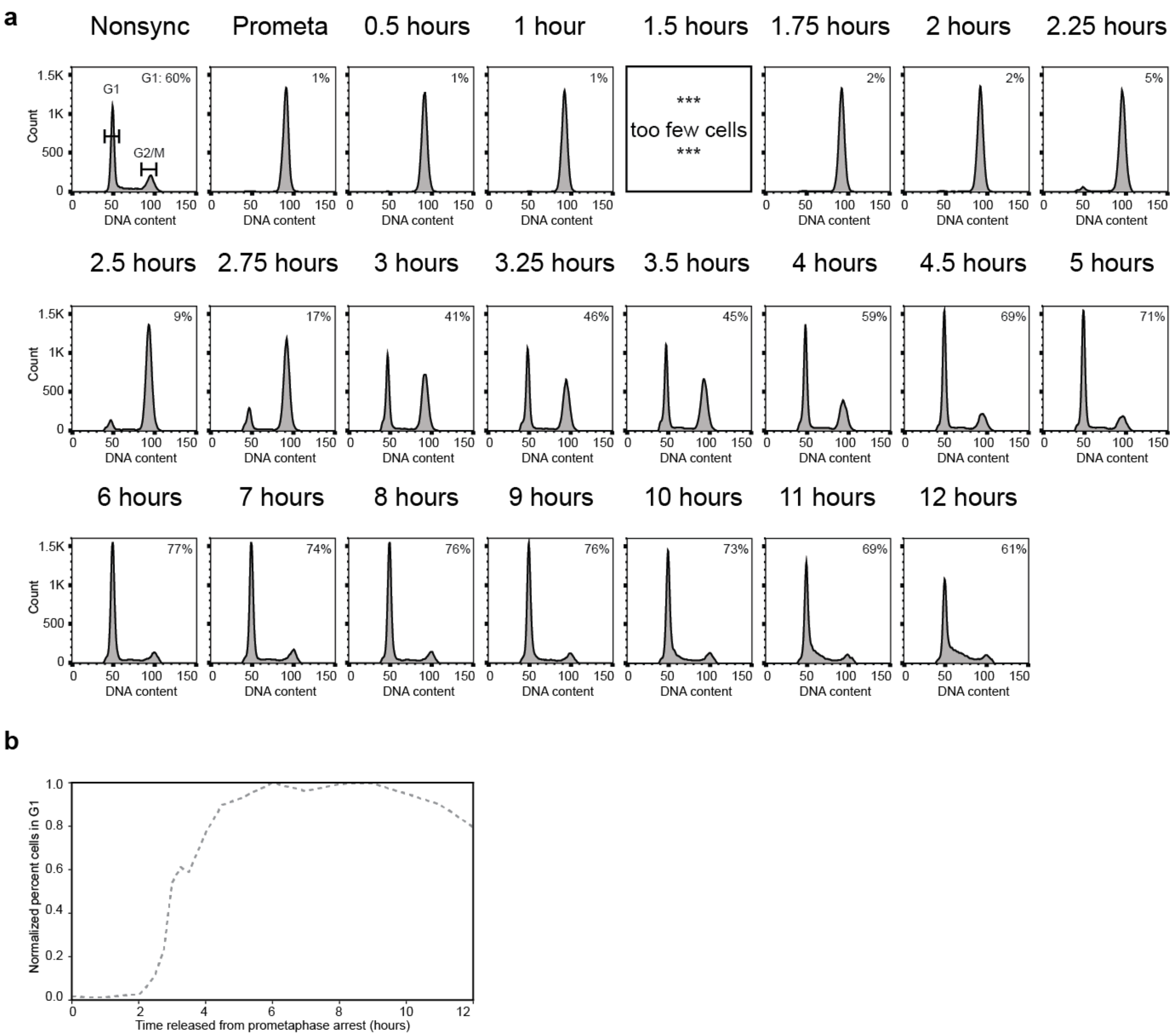


**Supplemental Fig. 1: Cell cycle analysis of mitotic exit for time course replicate 1**
**a,** FACS analysis of nonsynchronous and prometaphase-arrested cultures and of cultures at different time points after release from prometaphase-arrest. Percentages in the upper right corner represent the number of cells with a G1 DNA content. **b,** Quantification of the fraction of cells in G1 at each time point, normalized to t = 8 hours.

**a**

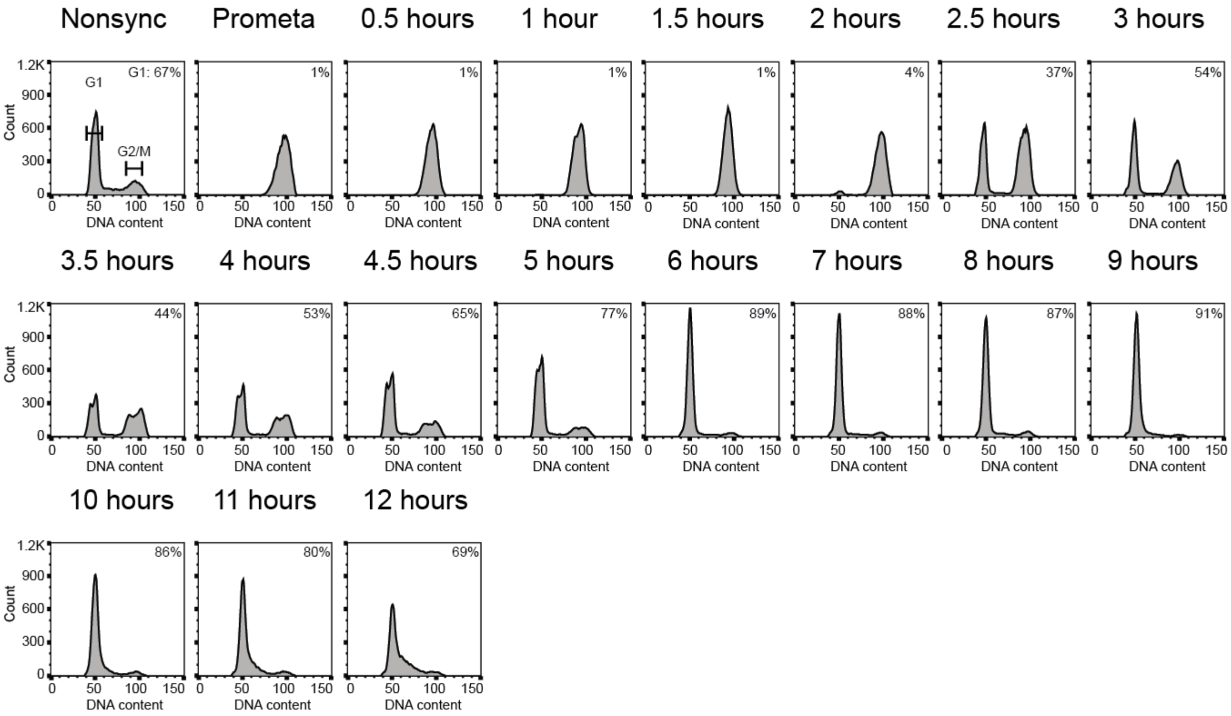


**b**

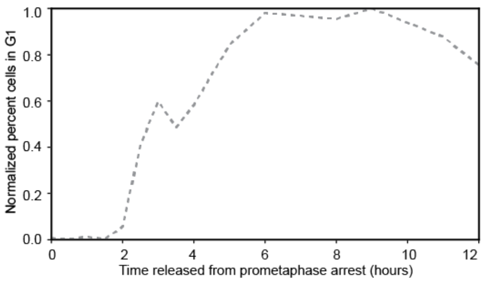


**Supplemental Fig. 2: Cell cycle analysis of mitotic exit for time course replicate 2**
**a,** FACS analysis of nonsynchronous and prometaphase-arrested cultures and of cultures at different time points after release from prometaphase-arrest. Percentages in the upper right corner represent the number of cells with a G1 DNA content. **b,** Quantification of the fraction of cells in G1 at each time point, normalized to t = 8 hours.

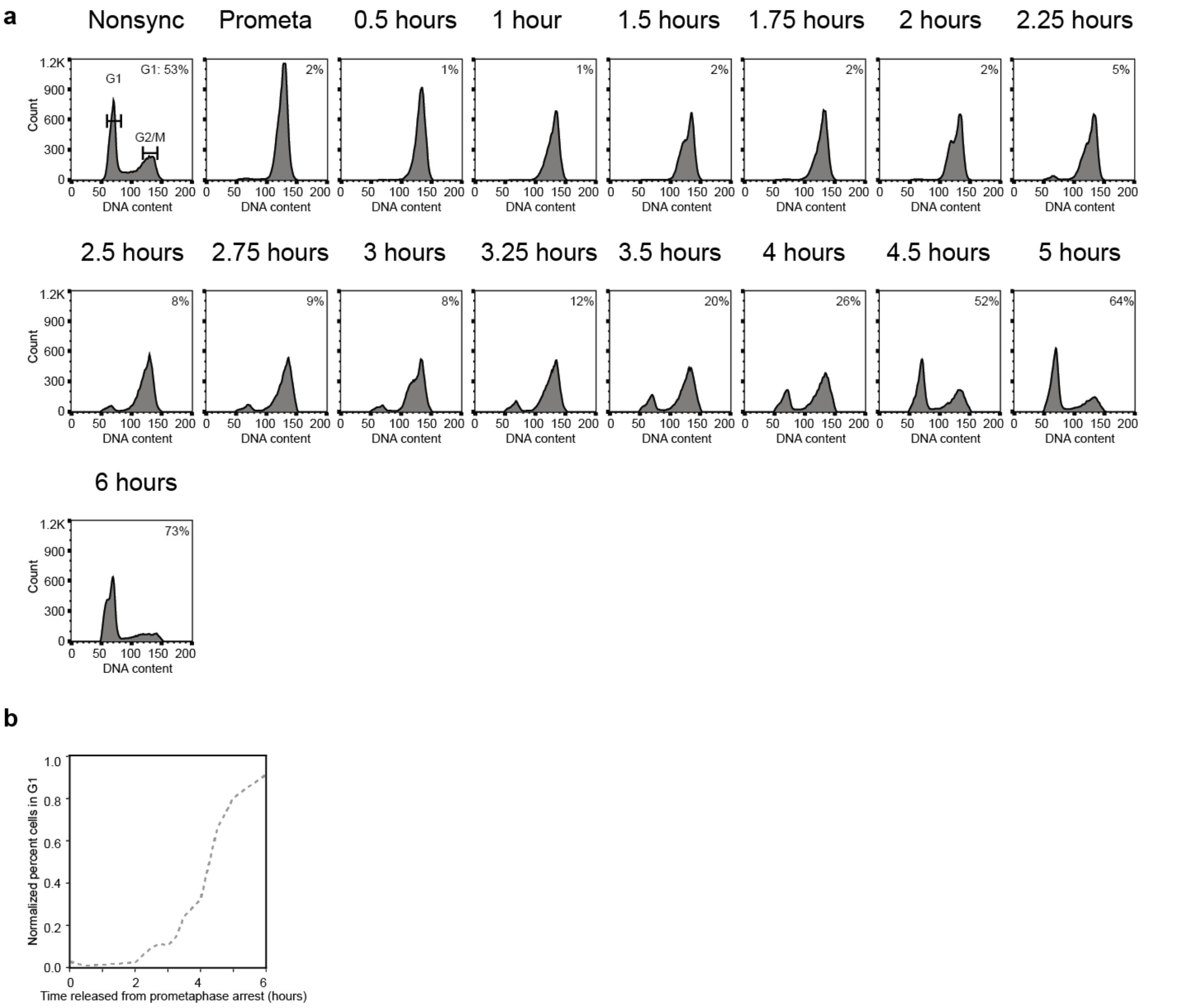


**Supplemental Fig. 3: Cell cycle analysis of mitotic exit for time course replicate 3**
**a,** FACS analysis of nonsynchronous and prometaphase-arrested cultures and of cultures at different time points after release from prometaphase-arrest. Percentages in the upper right corner represent the number of cells with a G1 DNA content. **b,** Quantification of the fraction of cells in G1 at each time point, normalized to G1 maximum assumed to be 80%.

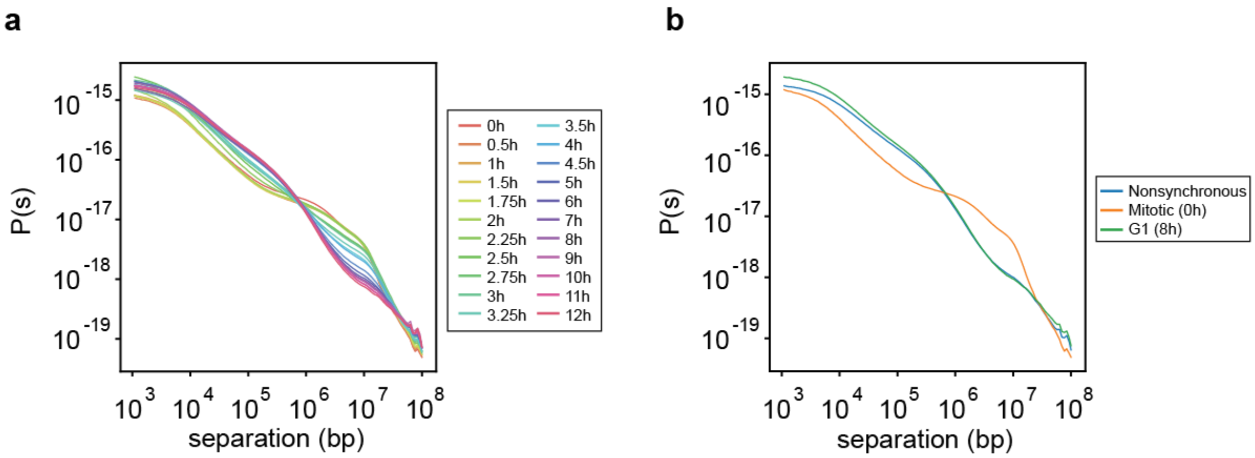


**Supplemental Fig. 4: *P*(*s*) plots from Hi-C data obtained with cells at different times after release from prometaphase**

**a,** *P*(*s*) plots for Hi-C data from cells at indicated time points after release from prometaphase.
**b,** *P*(*s*) plots for Hi-C data from nonsynchronous, mitotic (t = 0 hours), or G1 (t = 8 hours) cultures.

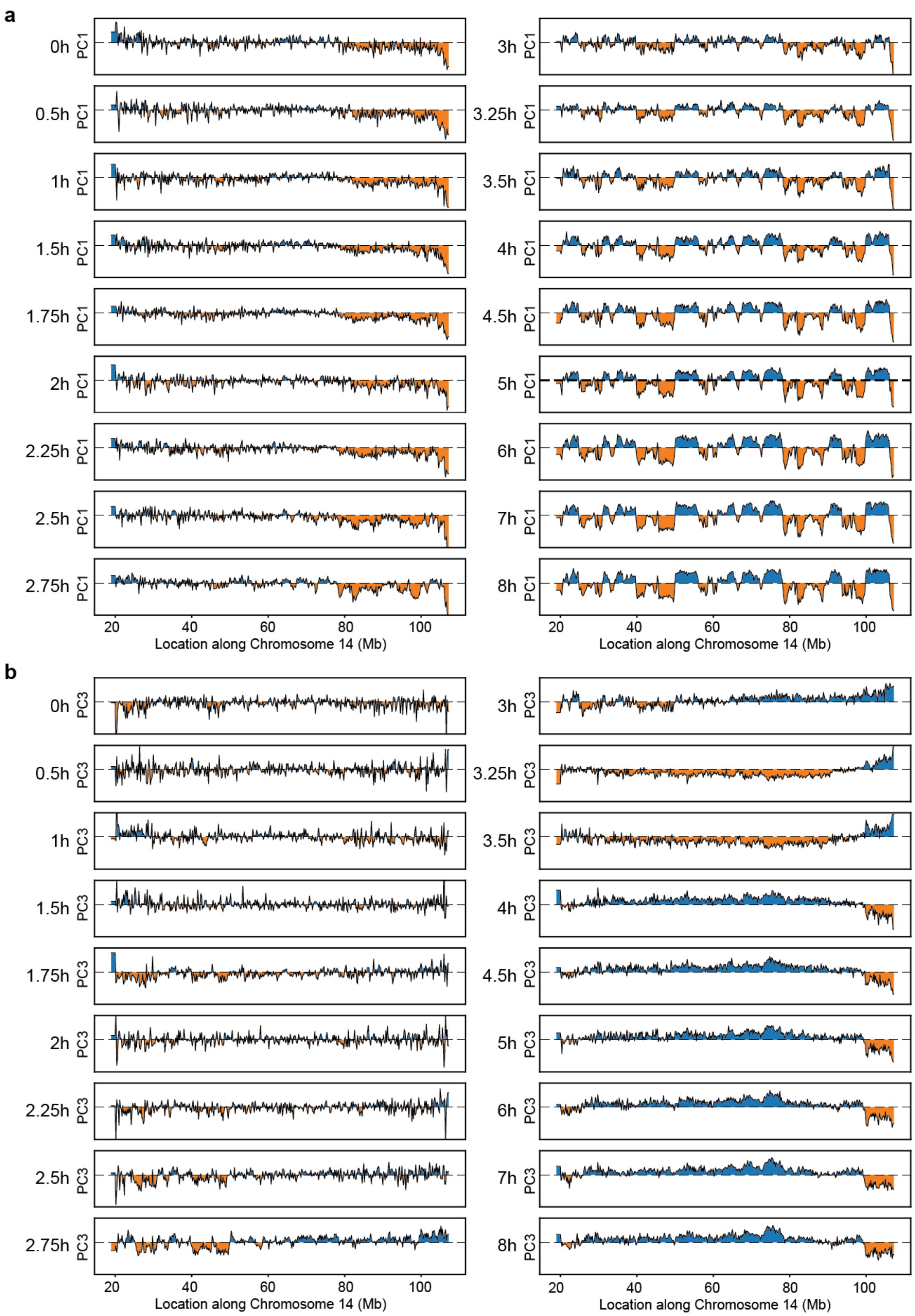
a
0h
0.5h
1h
1.5h
1.75h
2h
2.25h
2.5h
2.75h
3h
3.25h
3.5h
4h
4.5h
5h
6h
7h
8h
PC1
20
40
60
80
100
Location along Chromosome 14 (Mb)
b
PC3
Location along Chromosome 14 (Mb)

**Supplemental Fig. 5: Compartment analysis for time course replicate 1**
**a,** Principal component 1 (PC1) along Chromosome 14 for Hi-C data obtained from cells at different time points after release from prometaphase. Principal component analysis was performed on Hi-C data binned at 200 kb resolution. PC1 detects A and B compartments starting at t = 3 hours. **b,** Principal component 3 (PC3) along Chromosome 14 for Hi-C data obtained from cells at different time points after release from prometaphase. Principal component analysis was performed on Hi-C data binned at 200 kb resolution. PC3 detects some A and B compartments starting at t = 2.75 hours, but at later time points, PC1 captures compartments.

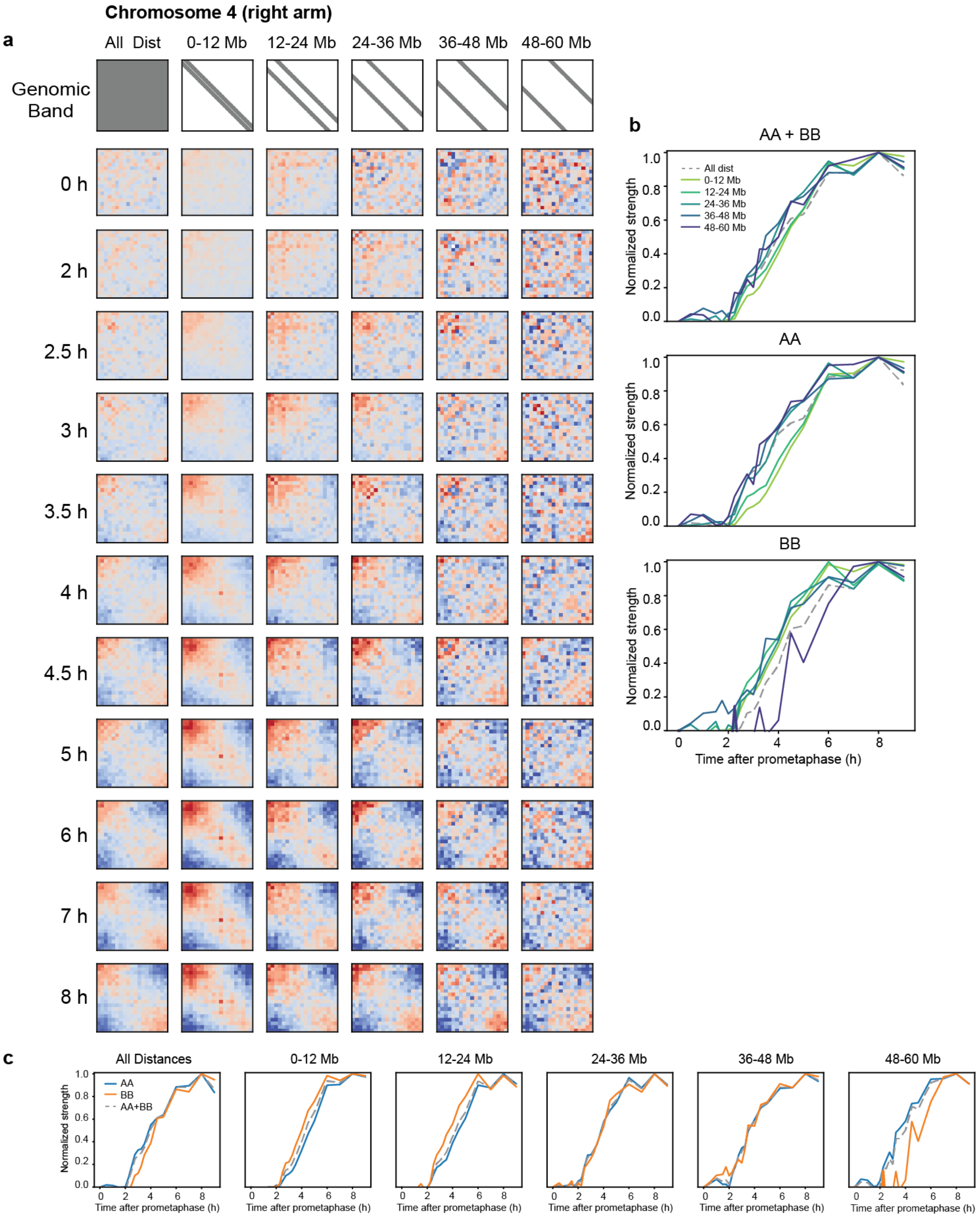
a
Chromosome 4 (right arm)
All Dist
0-12 Mb
12-24 Mb
24-36 Mb
36-48 Mb
48-60 Mb
Genomic Band
0 h
2 h
2.5 h
3 h
3.5 h
4 h
4.5 h
5 h
6 h
7 h
8 h
b
AA + BB
AA
BB
All dist
0-12 Mb
12-24 Mb
24-36 Mb
36-48 Mb
48-60 Mb
Normalized strength
Time after prometaphase (h)
c
All Distances
0-12 Mb
12-24 Mb
24-36 Mb
36-48 Mb
48-60 Mb
AA
BB
AA+BB
Normalized strength
Time after prometaphase (h)

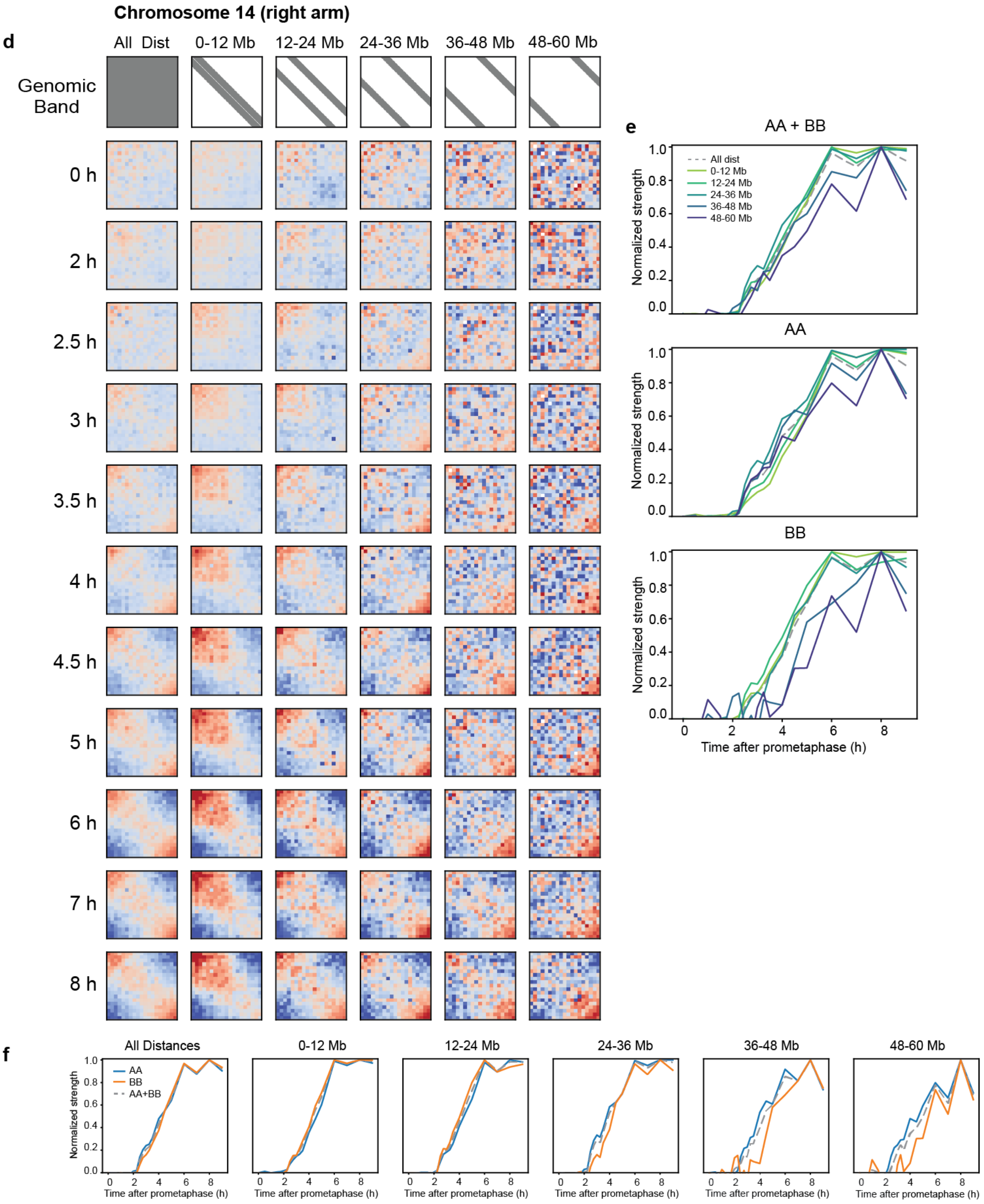

Chromosome 14 (right arm)
d
All Dist
0-12 Mb
12-24 Mb
24-36 Mb
36-48 Mb
48-60 Mb
Genomic Band
0 h
2 h
2.5 h
3 h
3.5 h
4 h
4.5 h
5 h
6 h
7 h
8 h
e
AA + BB
AA
BB
All dist
Normalized strength
Time after prometaphase (h)
f
All Distances
AA
BB
AA+BB

**Supplemental Fig 6: Compartment analysis at various genomic distances over time**
Saddle plots of Hi-C data binned at 200 kb resolution for different time points and split into genomic distance bands, as shown in gray in the first row, for chromosome 4 (**a**) and chromosome 14 (**d**). Normalized compartmentalization strength of different genomic distances as a function of time and split by interaction type (A-A, B-B, A-B) for chromosome 4 (**b**) and chromosome 14 (**e**). Normalized compartmentalization strength of interaction types as a function of time and split by genomic distance for chromosome 4 (**c**) and chromosome 14 (**f**).

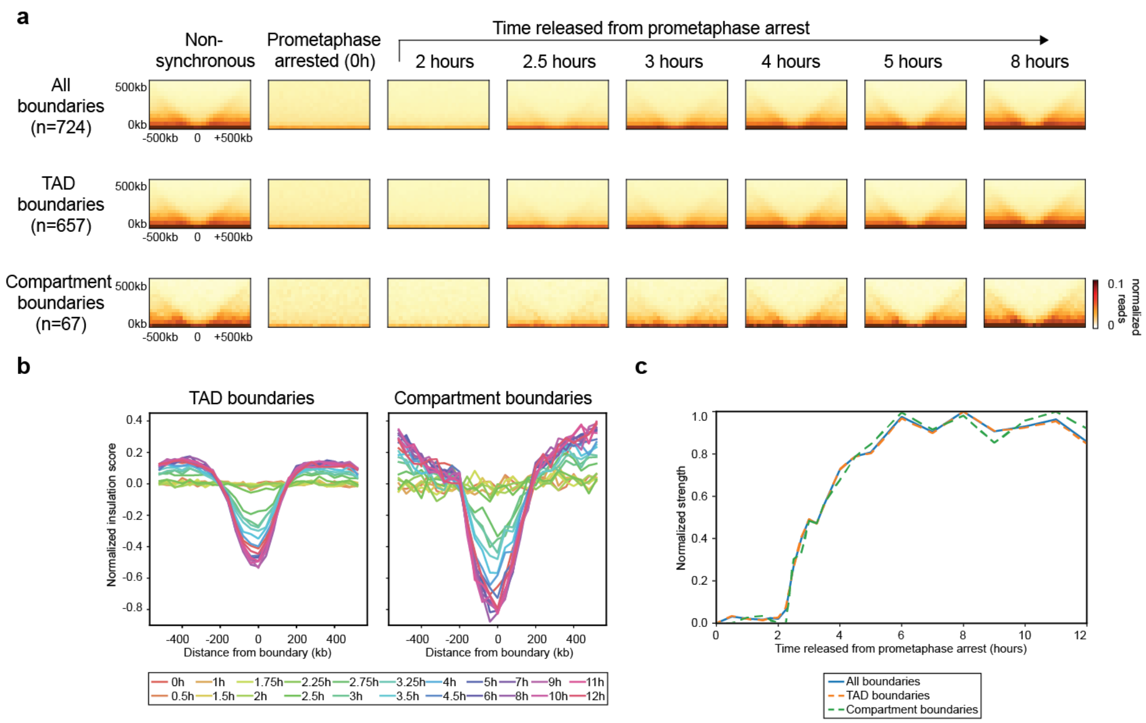


**Supplemental Fig. 7: TAD and compartment domain boundaries form with similar kinetics**

**a,** Aggregate Hi-C data binned at 40 kb resolution at domain boundaries (top = all boundaries, middle = TAD boundaries, bottom = compartment boundaries) at different time points after release from prometaphase. **b,** Average insulation profile across domain boundaries (left = TAD boundaries, right = compartment boundaries) for different time points. **c,** Normalized strength for domain boundaries as a function of time after release from prometaphase. The strength for each of these features was set at 1 for the 8 hour time point. TAD boundaries and compartment boundaries form with similar kinetics.

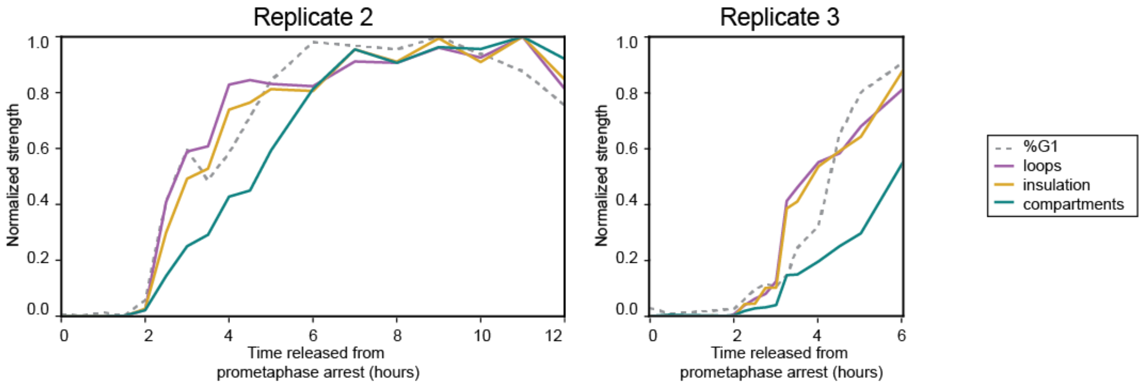


**Supplemental Fig. 8: Kinetics of chromosome feature formation in time course replicate 2 and 3**

Normalized feature strength for TADs, loops, and compartments as a function of time after release from prometaphase (Left: replicate time course 2, Right: replicate time course 3). For replicate 2: The strength for each of these features was set at 1 for the 8 hour time point. Dotted line indicates the fraction of cells in G1 at each time point, normalized to t = 8 hours. For replicate 3: The strength for each of these features was normalized to the strength expected based on data from replicate 1. Dotted line indicates the fraction of cells in G1 at each time point, normalized to G1 maximum assumed to be 80%.

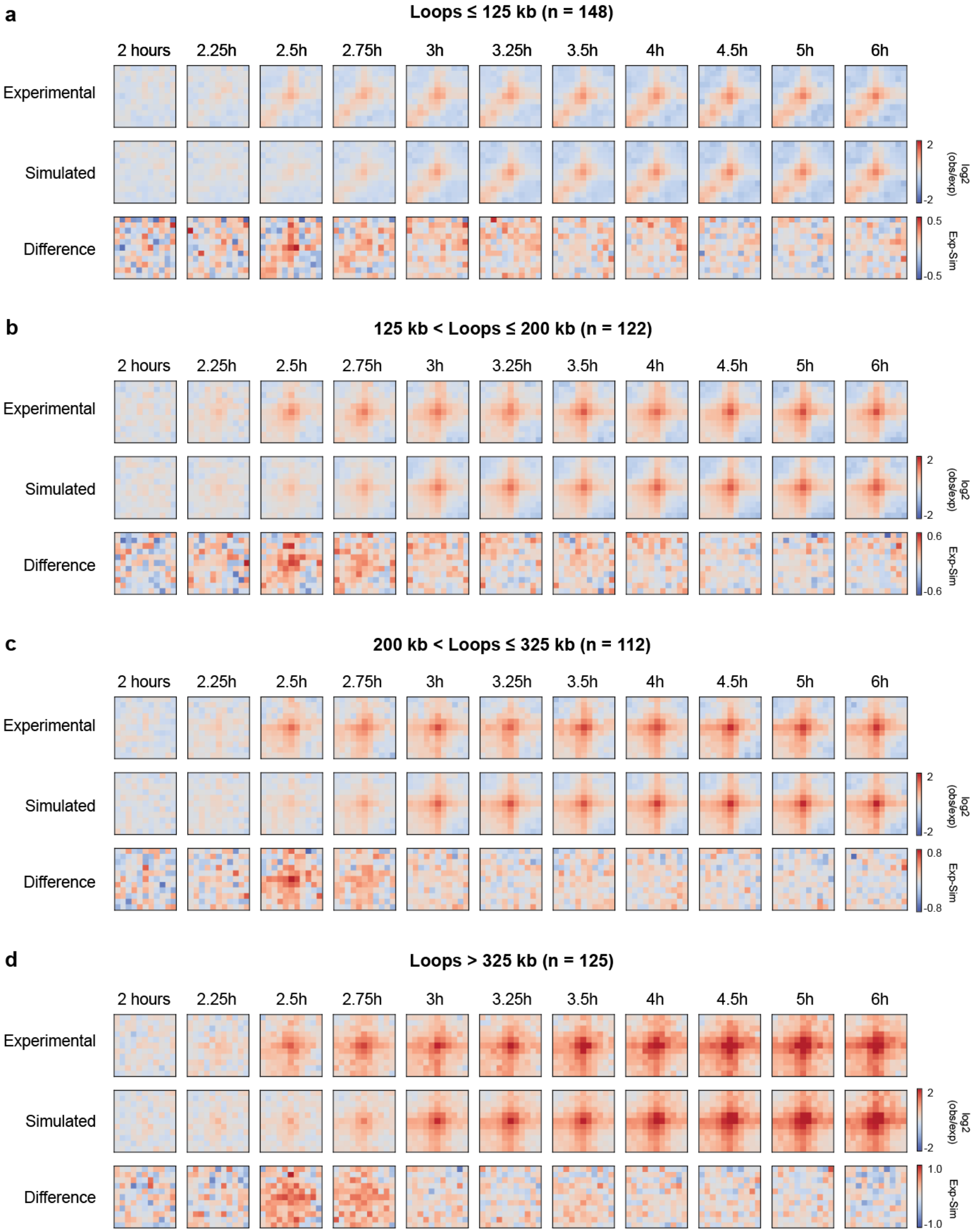
a
Loops ≤ 125 kb (n = 148)
b
125 kb < Loops ≤ 200 kb (n = 122)
c
200 kb < Loops ≤ 325 kb (n = 112)
d
Loops > 325 kb (n = 125)
2 hours
2.25h
2.5h
2.75h
3h
3.25h
3.5h
4h
4.5h
5h
6h
Experimental
Simulated
Difference
log2 (obs/exp)
Exp-Sim
2
-2
0.5
-0.5
0.6
-0.6
0.8
-0.8
1.0
-1.0

**Supplemental Fig. 9: Kinetics of loop formation for loops of different size**
Loops were grouped according to size: **a,** loops less than or equal to 125 kb, **b,** loops greater than 125 kb and less than or equal to 200 kb, **c,** loops greater than 200 kb and less than or equal to 325 kb, **d,** loops greater than 325 kb. For each panel, top row: log2(observed/expected) Hi-C data for experimental time course, middle row: log2(observed/expected) Hi-C data for simulated time course, bottom row: the difference between experimental and simulated Hi-C data. Kinetics of loop formation is similar for all loop sizes.

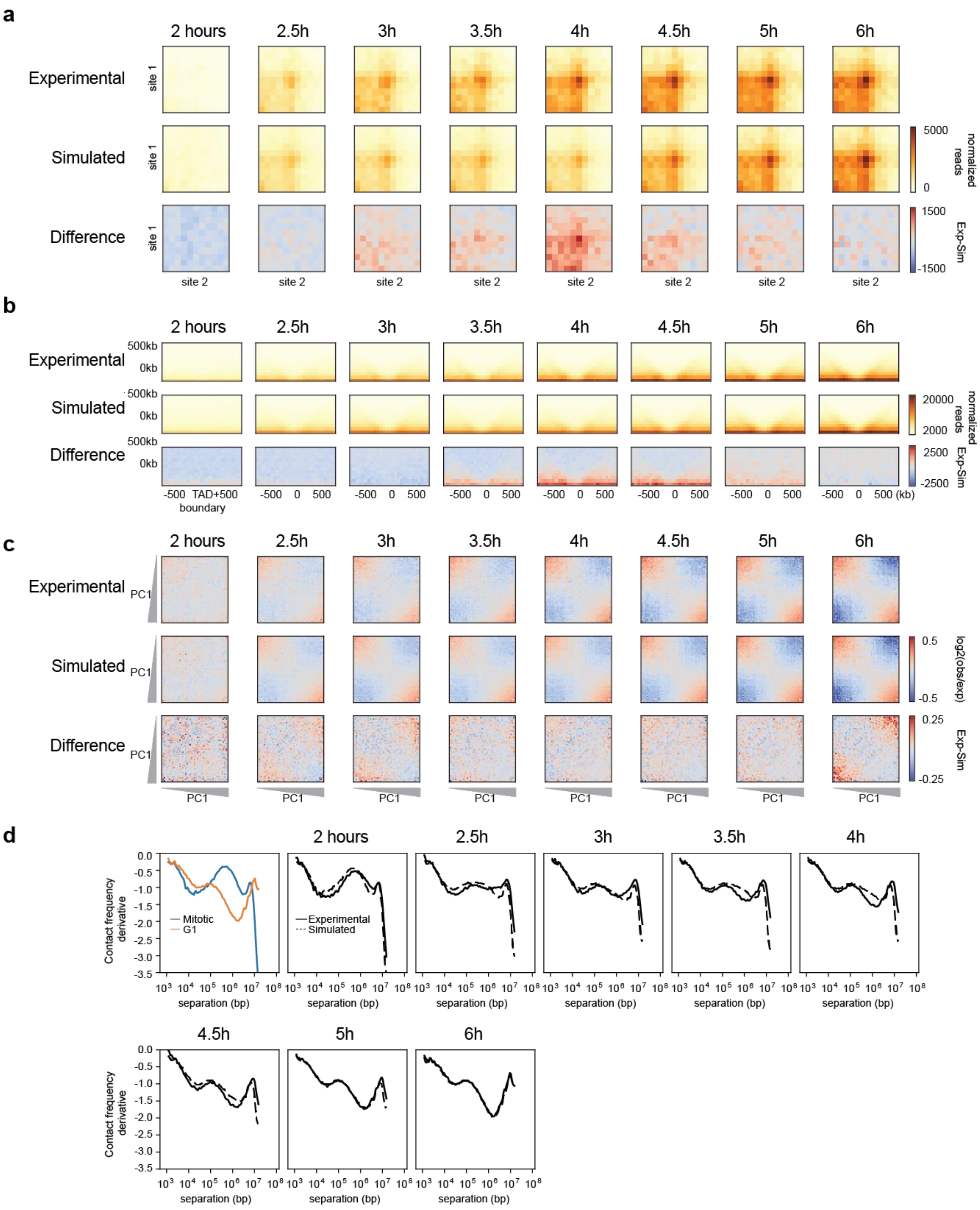

a
2 hours
2.5h
3h
3.5h
4h
4.5h
5h
6h
Experimental
Simulated
Difference
site 1
site 2
5000
0
normalized reads
1500
-1500
Exp-Sim
b
500kb
0kb
20000
2000
2500
-2500
-500 TAD+500 boundary
-500 0 500
500 (kb)
c
PC1
0.5
-0.5
log2(obs/exp)
0.25
-0.25
d
Contact frequency derivative
Mitotic
G1
Experimental
Simulated
separation (bp)

**Supplemental Fig. 10: Analysis of time course replicate 2**

**a,** Aggregated Hi-C data binned at 20 kb resolution at chromatin loops at different time points. Top row: Experimental Hi-C data. Middle row: Simulated Hi-C data. Bottom row: The difference between experimental and simulated Hi-C data. Loops are more prominent in experimental Hi-C data than in the simulated data between 3 and 4.5 hours. This analysis included loops larger than 200 kb to avoid the strong signal at the diagonal of the interaction matrix. Simulations were performed with experimental data from this time course (mixing Hi-C data for t = 0 and t = 8 hours). **b,** Aggregate Hi-C data binned at 40 kb resolution at TAD boundaries for different time points. Top row: Experimental Hi-C data. Middle row: Simulated Hi-C data. Bottom row: The difference between experimental and simulated Hi-C data. Insulation strength is stronger in experimental Hi-C data than in simulated Hi-C data at t = 3.5 and t = 4.5 hours. **c,** Saddle plots of Hi-C data binned at 200 kb resolution for different time points. Top row: Experimental Hi-C data. Middle row: Simulated Hi-C data. Bottom row: The difference between experimental and simulated Hi-C data. Compartmentalization is weaker in experimental Hi-C than in simulated Hi-C data as illustrated by the fact that A-B interactions are less depleted in the experimental data (upper right and lower left corner of saddle plots). **d,** Derivative from $P(s)$ plots. Solid lines represent the derivative of $P(s)$ for experimental Hi-C data and the dotted lines represent the derivative of $P(s)$ for the simulated Hi-C datasets for corresponding time points.

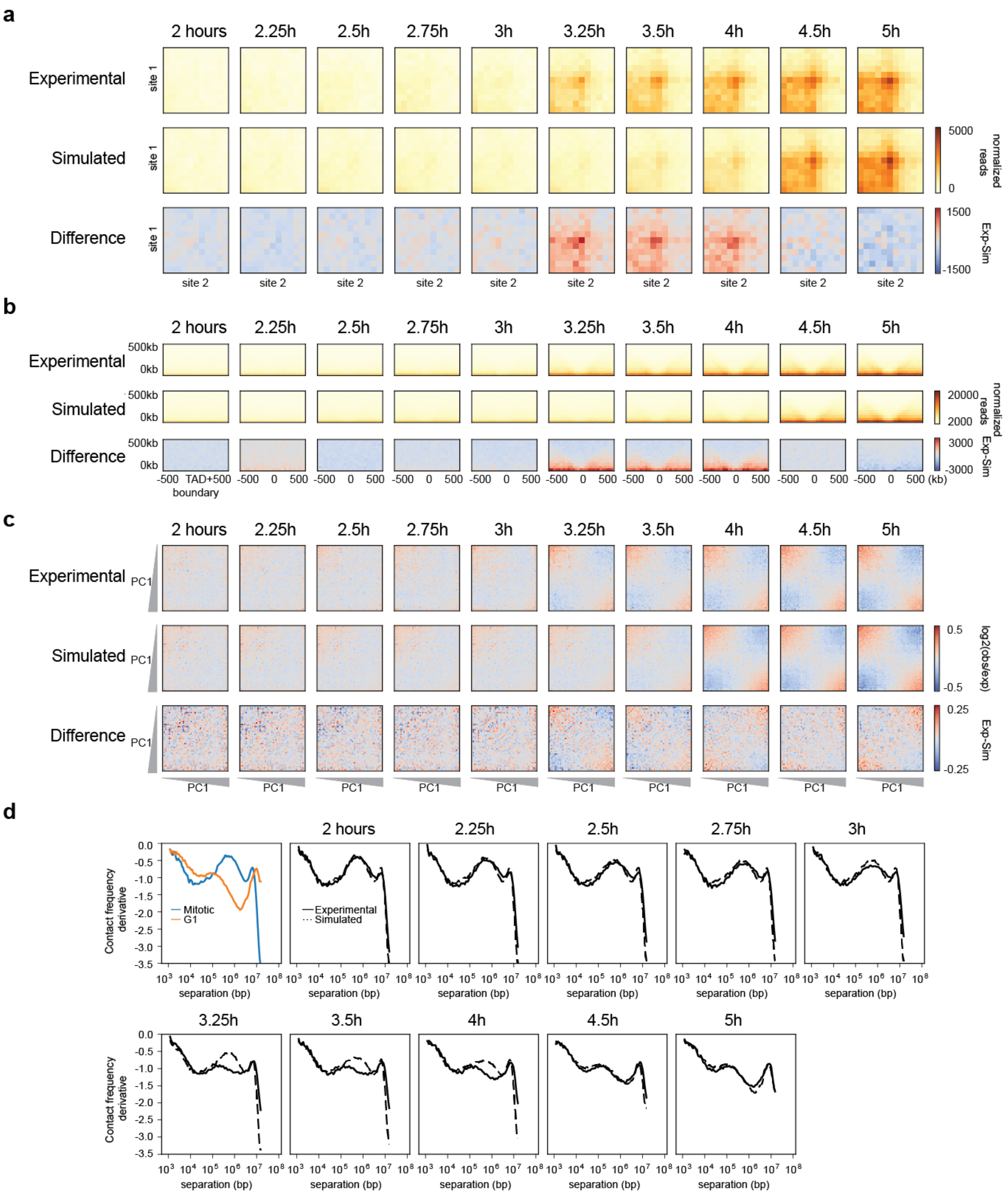
a
2 hours
2.25h
2.5h
2.75h
3h
3.25h
3.5h
4h
4.5h
5h
Experimental
Simulated
Difference
site 1
site 2
5000
0
normalized reads
1500
-1500
Exp-Sim
b
500kb
0kb
-500 TAD+500 boundary
-500 0 500
(kb)
20000
2000
3000
-3000
c
PC1
0.5
-0.5
log2(obs/exp)
0.25
-0.25
d
Contact frequency derivative
separation (bp)
Mitotic
G1
Experimental
Simulated

**Supplemental Fig. 11: Analysis of time course replicate 3**

**a,** Aggregated Hi-C data binned at 20 kb resolution at chromatin loops at different time points. Top row: Experimental Hi-C data. Middle row: Simulated Hi-C data. Bottom row: The difference between experimental and simulated Hi-C data. Loops are more prominent in experimental Hi-C data than in the simulated data between 3.25 and 4 hours. This analysis included loops larger than 200 kb to avoid the strong signal at the diagonal of the interaction matrix. Simulations were performed with experimental data from this time course (mixing Hi-C data for t = 0 and t = 6 hours). **b,** Aggregate Hi-C data binned at 40 kb resolution at TAD boundaries for different time points. Top row: Experimental Hi-C data. Middle row: Simulated Hi-C data. Bottom row: The difference between experimental and simulated Hi-C data. Insulation strength is stronger in experimental Hi-C data than in simulated Hi-C data at t = 3.25 and t = 4 hours. **c,** Saddle plots of Hi-C data binned at 200 kb resolution for different time points. Top row: Experimental Hi-C data. Middle row: Simulated Hi-C data. Bottom row: The difference between experimental and simulated Hi-C data. Compartmentalization is weaker in experimental Hi-C than in simulated Hi-C data as illustrated by the fact that A-B interactions are less depleted in the experimental data (upper right and lower left corner of saddle plots). **d,** Derivative from *P*(*s*) plots. Solid lines represent the derivative of *P*(*s*) for experimental Hi-C data and the dotted lines represent the derivative of *P*(*s*) for the simulated Hi-C datasets for corresponding time points.

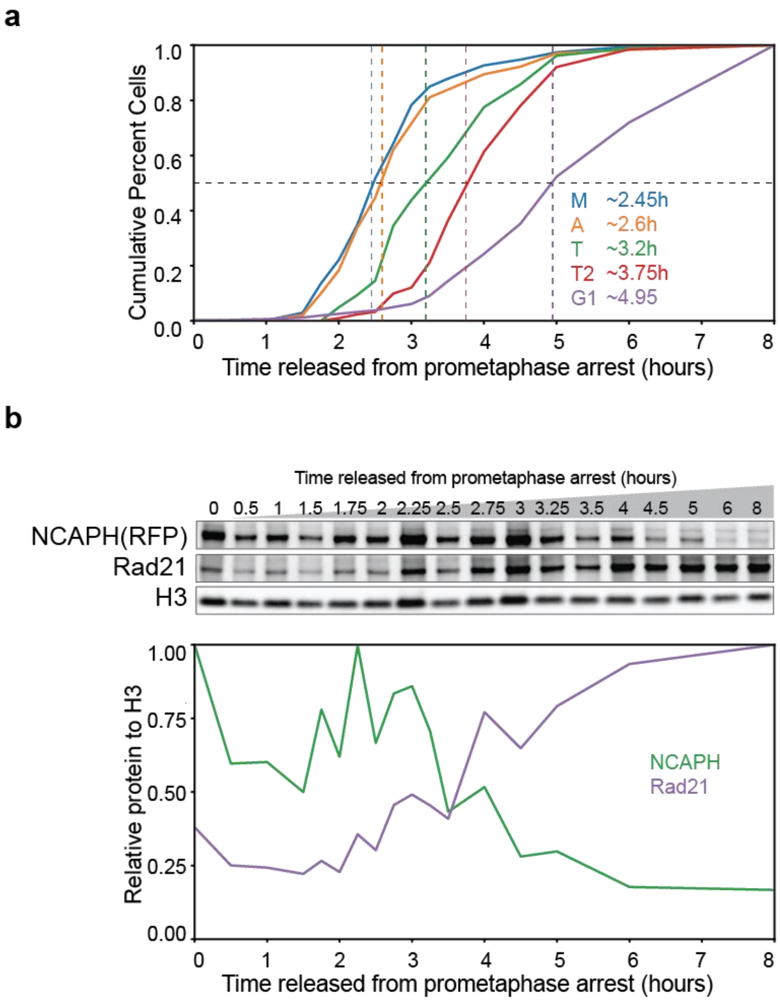


**Supplemental Fig. 12: Mitotic exit kinetics and chromatin association dynamics of condensin and cohesin for HeLaS3-NCAPH-dTomato cells**

**a,** Cumulative plots of HeLaS3-NCAPH-dTomato cells at different cell cycle stages defined by imaging. Classification of cell cycle stages was based on DAPI staining and alpha-tubulin organization. Prometaphase cells were defined as cells with condensed chromosomes and disrupted tubulin structure due to the microtubule inhibitor used for prometaphase arrest. Cells classified as metaphase had a single axis of DAPI staining with tubulin aligned on each side. Anaphase cells had tubulin on each side of the DAPI axis but must have had two distinct DAPI clusters representing the separation of two genomic copies. Late anaphase/early telophase classification was characterized by the presence of tubulin only between the two DAPI populations and no longer on the ends. When the tubulin signal was compressed between the two DAPI clusters, we classified those as late telophase and cytokinesis. All cells with decondensed chromatin and no nuclear tubulin were classified as G1 cells. M = metaphase, A = early anaphase, T = late anaphase/early telophase, T2 = late telophase, G1 = G1. **b,** Top: Western blot analysis of chromatin-associated proteins purified from cells at different time points after release from prometaphase. Bottom: Quantification of the western blot shown above. NCAPH and Rad21 were analyzed on the same gel. The samples for Histone H3 analysis were run on another gel.
